# An Optimized Product-Enhanced Reverse Transcriptase Assay for Sensitive and Quantitative Detection of HIV Viral Load and Phenotypic Drug Resistance

**DOI:** 10.64898/2025.12.10.691922

**Authors:** Dorothy K. Mims, Megan M. Chang, Ayokunle O. Olanrewaju

**Affiliations:** Department of Bioengineering, University of Washington, Seattle, WA, 98105; Department of Mechanical Engineering, University of Washington, Seattle, WA, 98105

**Author notes:** Corresponding Author: Ayokunle Olanrewaju.

## Abstract

The World Health Organization (WHO) recommends Human Immunodeficiency Virus (HIV) drug resistance testing (DRT), but current tests are too complex and expensive for routine use. Genotypic DRT is challenging to interpret because of the growing list of mutations responsible for HIV drug resistance. Although phenotypic DRT is simpler to interpret, it requires slow and labor-intensive viral culture. Phenotypic tests that measure the activity of isolated HIV enzymes are faster and less labor-intensive, but none have yet met the 2023 WHO Target Product Profile (TPP) for HIV DRT. Here we present an optimized Product-Enhanced Reverse Transcriptase (PERT) assay for sensitive and quantitative detection of HIV viral load and drug resistance.

PERT combines complementary DNA (cDNA) synthesis by HIV-Reverse Transcriptase (HIV-RT) with cDNA amplification and detection by quantitative PCR (qPCR). We established sensitive detection down to 10 HIV-RT molecules in a 25 µL sample corresponding to a viral load of ∼10 HIV RNA copies/mL. We demonstrated the assay’s feasibility for phenotypic DRT using lamivudine-5’-triphosphate (3TC-TP)—to which the M184V mutation confers high-level resistance—and quantified 3TC-TP inhibition using the difference in cDNA produced between drug and no-drug conditions. We surpassed WHO minimal analytical sensitivity requirements (20% low-abundance variant detection) by differentiating 10% M184V HIV-RT fractions in heterogenous mixtures (1,000 total HIV-RT, viral load ∼2,500 copies RNA/mL). Finally, we showed that PERT could detect resistance to existing and emerging RT inhibitors including tenofovir-diphosphate (TFV-DP), doravirine (DOR), and islatravir (ISL)-TP.

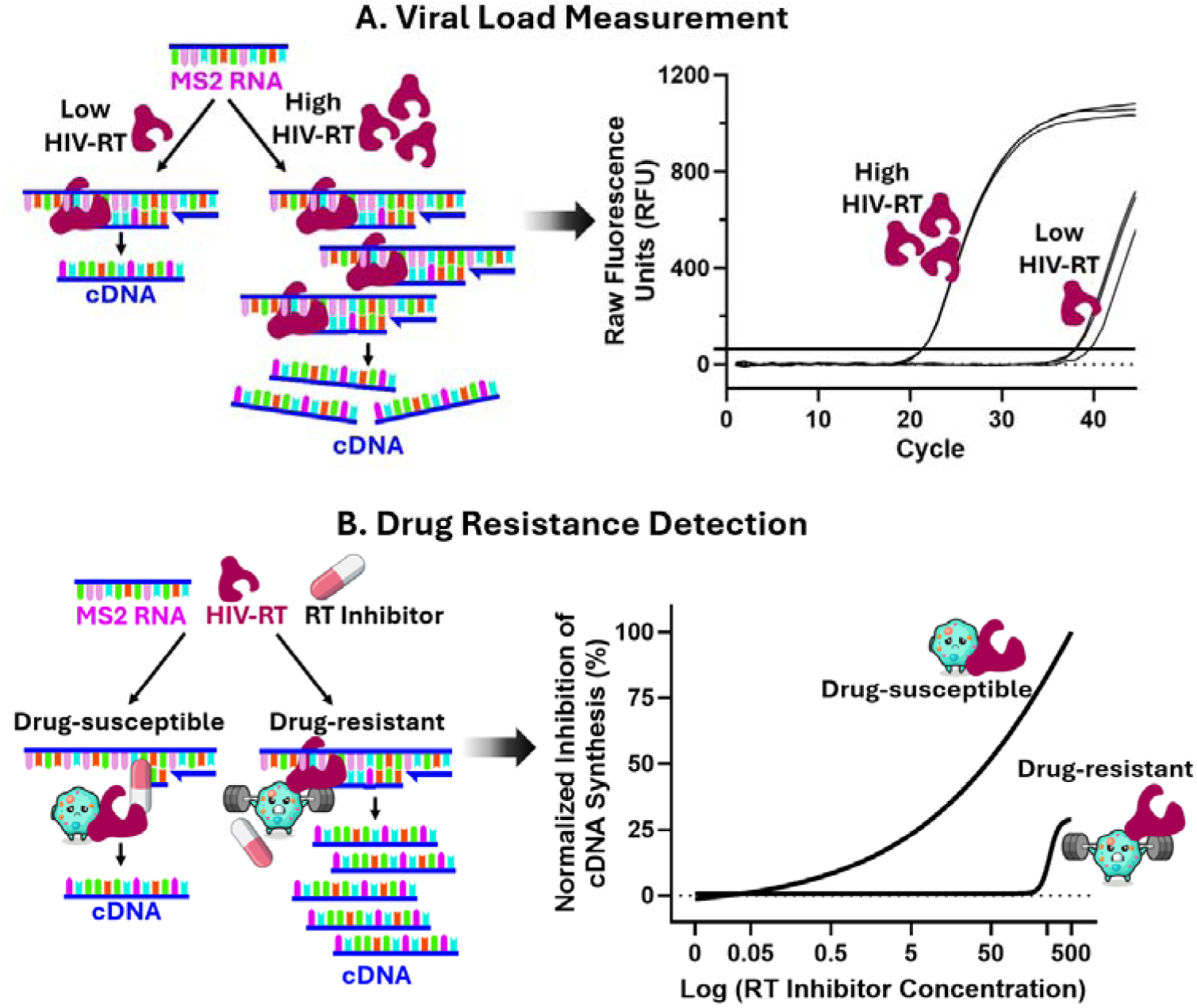

The Product-Enhanced Reverse Transcriptase (PERT) assay measures HIV-RT activity by its ability to synthesize complementary DNA, providing a phenotypic measurement of HIV viral load (by quantity of HIV-RT enzyme) and susceptibility to RT inhibitors (by level of inhibition of HIV-RT activity).

**IMPORTANCE:** Although antiretroviral therapy can effectively treat and prevent HIV infection, treatment efficacy and global control of the HIV epidemic are threatened by rising rates of HIV drug resistance. Inexpensive and decentralized HIV DRT could facilitate surveillance efforts to understand the prevalence of HIV drug resistance in local and global contexts. This need is especially timely and pressing considering anticipated increased rates of HIV acquisition and drug resistance due to the reductions in global HIV services driven by recent funding cuts. Our optimized PERT assay for simultaneous viral load and phenotypic drug resistance quantification is fast (∼2 hours), sensitive, and accurate. We can also leverage existing RT-qPCR instruments used for viral load measurement to significantly improve access to HIV drug resistance monitoring and tailored regimen selection.

## INTRODUCTION

Human Immunodeficiency Virus (HIV) affects more than 39 million people worldwide, with over 600,000 HIV-related deaths in 2023 alone.^1^ People living with HIV (PLWH) receiving antiretroviral therapy (ART) can achieve viral suppression and undetectable viral load (VL), which leads to drastically increased life expectancy and no risk of transmission.^2^ Despite 30.7 million people accessing ART, many PLWH do not achieve viral suppression—only 65% of PLWH in the U.S. and 72% globally were virally suppressed as of 2023.^1^ Virologic failure (> 200 copies/mL viral RNA^3,4^) on ART typically arises due to either medication non-adherence or drug resistance; insufficient medication adherence results in low drug levels that can lead to virologic failure, creating ideal conditions for the emergence of drug resistance. In some regions with high HIV prevalence, drug resistance rates are concerningly high^5^: 75% for any class of ART, 66% for non-nucleoside reverse transcriptase inhibitors (NNRTIs), 42% for nucleoside reverse transcriptase inhibitors (NRTIs), and 16% for integrase strand transfer inhibitors (INSTIs)—a drug class approved less than a decade ago.

Although the World Health Organization (WHO) recommends resistance monitoring as part of HIV treatment and prevention programs,^6^ this is not typically done in practice—particularly in low-and middle-income countries (LMICs)—because of the associated cost and complexity. In some high HIV prevalence countries (e.g. South Africa), drug resistance testing (DRT) is typically delayed or rationed given the high test cost—instead, PLWH are often “blindly switched” to more expensive second-line regimens.^7^ Delayed DRT and regimen switches can provide additional time for new mutations to develop and for drug resistance to be transmitted to other people, jeopardizing individual- and population-level control of the HIV epidemic.

There are two main types of DRT: genotypic and phenotypic. Genotypic DRT includes point-mutation assays,^8^ DNA amplification-based assays that analyze known resistance-associated mutations,^9–11^ and sequencing methods including Sanger sequencing and next-generation sequencing.^12^ While highly sensitive, point-mutation assays and DNA amplification-based methods can only determine resistance due to a limited number of known mutations and can miss emerging yet clinically relevant mutations.^13^ Sequencing-based methods address this concern, but are more expensive, labor-intensive, and challenging to interpret. Phenotypic DRT measures HIV replication in the presence of antiretroviral drugs, and does not require prior knowledge of the mutations. However, phenotypic DRT used in reference labs, like the PhenoSense® assay (Monogram Biosciences),^14^ requires labor-intensive and slow viral culture by highly trained operators. Phenotypic DRT that isolates and tests the activity of an HIV enzyme in the presence of antiretroviral drugs^15^ does not require viral culture and offers potential for more widespread application outside centralized labs. These phenotypic enzymatic methods have been in development for over 30 years^16^ and can estimate both HIV VL^17–20^ and drug resistance.^17,21,22^ For example, the Product-Enhanced Reverse Transcriptase (PERT) assay measures RT activity by combining complementary DNA (cDNA) synthesis by an RT enzyme with amplification and detection by quantitative PCR (qPCR).

Phenotypic enzymatic assays like PERT are relatively less explored for HIV diagnostic testing than culture-based phenotypic assays and genotypic/sequencing methods. Although phenotypic assays are generally expected to cover all mutations and subtypes, most fall short in sensitivity, with no phenotypic enzyme activity assays yet meeting the sensitivity guidelines provided in the 2023 WHO Target Product Profile (TPP) for HIV DRT. The WHO TPP lists requirements for HIV DRT in LMICs regarding the scope; assay design, performance, and functionality; specimen handling; report and data handling; cost considerations; and extraneous requirements. In this manuscript we aimed to address the main sensitivity requirements within assay design, performance, and functionality, which we believe to be the first steps in meeting the TTP guidelines and which previous phenotypic enzymatic assays have been unable to meet. Specifically, the minimal WHO TPP sensitivity requirements for HIV DRT in LMICs are detecting 20% drug resistant fraction at a level of viremia ranging from 1,000–5,000 copies RNA/mL.^15^

Here we present an optimized PERT assay with improved turnaround time and increased analytical sensitivity, detecting as low as 10 molecules of wildtype (WT) or mutant HIV-RT enzyme. We then demonstrated that a modified buffer could support extraction-free testing by direct lysis of HIV virions and addition to downstream cDNA synthesis and qPCR reactions with minimal inhibition by the presence of plasma. As a proof of concept for HIV DRT, we compared WT HIV-RT with the M184V mutation, which confers high-level resistance to the NRTI drug lamivudine-5’-triphosphate (3TC-TP) that is used in WHO-recommended first line ART regimens.^23,24^ We distinguished mixtures containing 10% fraction of the M184V mutant from WT HIV-RT at 500 µM 3TC-TP, exceeding the sensitivity requirements in the WHO TPP. We also showed distinction of WT from M184V, K65R, and Y188L HIV-RT using the nucleoside reverse transcriptase translocation inhibitor (NRTTI) islatravir (ISL)-TP, the NRTI tenofovir-diphosphate (TFV-DP), and the NNRTI doravirine (DOR), respectively. This highly sensitive assay takes ∼2 hours to complete and uses existing RT-qPCR instruments that are available in settings where HIV VL can be measured, with the potential to significantly improve access and reduce instrumentation costs required for HIV DRT compared with genotypic and phenotypic culture-based methods.

## MATERIALS AND METHODS

### Buffer, Enzyme, RNA, Plasma, and Drug Handling and Storage

#### HIV-Reverse Transcriptase Buffer

1X HIV-Reverse Transcriptase (HIV-RT) Buffer was prepared containing 50mM Tris-HCl (pH 8.0, Invitrogen; Carlsbad, California, USA; 15568-025), 50 mM Potassium Chloride (Sigma-Aldrich; St. Louis, Missouri, USA; 60142- 100ML-F), 10mM MgCl2 (Sigma-Aldrich; St. Louis, Missouri, USA; M1028), 5 mM Dithiothreitol (DTT, Thermo Fisher Scientific™; Waltham, Massachusetts, USA; R0861), 0.16% Triton X-100 (Sigma-Aldrich; St. Louis, Missouri, USA; 9036-19-5), and nuclease-free (NF) water (Invitrogen; Carlsbad, California, USA; AM9937). RT Buffer was prepared in 5X and 7.5X mixtures and stored at - 20°C.

#### HIV-RT Buffer for Direct Plasma and HIV-1 Cell Culture Supernatant Testing

1X RT Buffer was prepared using higher Triton X (1.6% Triton X in 1X RT Buffer) and reduced DTT (2.5 mM DTT in 1X RT Buffer) for use with plasma and culture supernatant.

#### HIV-Reverse Transcriptase (HIV-RT) Source, Dilution, and Storage

Recombinant wildtype (WT) HIV-RT was acquired from Worthington Biochemical Corporation (Lakewood, NJ, USA; LS05009). The recombinant M184V Mutant HIV-RT was obtained through the NIH HIV Reagent Program, Division of AIDS, NIAID, NIH: Human Immunodeficiency Virus Type 1 HXB2 Reverse Transcriptase/M184V Heterodimeric Protein, Recombinant from Escherichia coli, ARP- 3195, contributed by Dr. Vinayaka Prasad and Dr. Mark Wainberg (BEI Resources, ARP-3195). Additional recombinant M184V as well as K65R and Y188L HIV-RT were kindly provided by Dr. Barry Lutz and Enos Kline.

Upon receipt, HIV-RT was divided into 0.5 to 1 µL aliquots (without dilution) and stored at - 20°C (WT) or -80°C (M184V) until dilution in 1X HIV-RT Buffer. For use in the assay, HIV-RT was diluted to the appropriate concentration (molecules/µL) in 1X HIV-RT Buffer. The manufacturer’s reported stock concentration (19.9 U/µL) and minimum activity (5,000 U/mg) were used to calculate the concentration of HIV-RT in µg/µL, and the molecular weight of HIV-RT (117,000 g/mol) and Avogadro’s number (6.022 E+23 molecules/mol) used to further derive the concentration in molecules HIV-RT/µL. For expressed mutant HIV-RT, concentration (µg/µL) was measured using a NanoDrop spectrophotometer and verified by standard curve alongside recombinant WT HIV-RT; measured concentration was then converted to HIV-RT molecules/µL as explained above. Once in RT Buffer, all aliquots (WT or mutant) were kept at -80°C to preserve the enzyme from degradation; aliquot volumes were chosen to minimize freeze-thaw cycles of enzyme after the initial dilution. All aliquots of HIV-RT were kept in Protein LoBind™ tubes (Eppendorf; Hamburg, Germany; 022431081 or 022431064).

HIV-1 cell culture supernatant was obtained from Dr. Geoffrey S. Gottlieb and Robert A. Smith: HIV-1 NL4-3 was grown for 8 days in CEM-SS cells and infectious titer verified by MAGIC-5A assay to be 2,713,600 focus-forming units (FFU). Culture supernatant was stored at -80°C until use then thawed and diluted using the modified 1X RT Buffer made with 1.6% Triton X and 2.5 mM DTT.

#### MS2 Bacteriophage RNA

(Millipore Sigma; Burlington, Massachusetts, USA; 10165948001) was divided into aliquots in 0.5 µL tubes upon receipt and stored at -20°C.

**Lamivudine-5’-triphosphate (3TC-TP)** was synthesized by Sierra Bioresearch (Tucson, Arizona, USA; Lot No. LM2558) and diluted in nuclease-free water then stored at -20°C. **Islatravir (ISL)-TP** (MedChemExpress, Monmouth Junction, New Jersey, USA; HY-138561) was reconstituted in nuclease-free water and stored at -20°C. **Tenofovir-diphosphate (TFV-DP)** (LGC Standards, Teddington, Middlesex, UK; TRC-T018525) was reconstituted in nuclease-free water and stored at -20°C. **Doravirine (DOR) (**BOC Bioscience, Shirley, New York, USA; B2693-475314) was reconstituted in dimethyl sulfoxide (DMSO) (Fisher Scientific; Waltham, Massachusetts; BP231-100) and stored in 50 mM aliquots at 4°C then further diluted in nuclease-free water and stored at -20°C.

#### Plasma preparation and storage

Verified non-viral pooled donor blood was obtained from BioIVT (Woodbury, New York, USA) and plasma prepared by centrifuging blood (upon receipt) at 2000 g for 10 minutes in a refrigerated centrifuge (4°C, in EDTA-coated tubes) then transferring plasma to Protein LoBind™ tubes and storing at -20°C or -80°C until use.

### Primers and Probe for HIV-RT Activity Assay

The forward primer (5’ - AACATGCTCGAGGGCCTTA - 3’), reverse primer (5’- GCCTTAGCAGTGCCCTGTCT - 3’), and probe (PrimeTime™ qPCR Probe: 5’ – FAM – CCCGTGGGATGCTCCTACATGTCA – TAMRA – 3’) as designed by Ma and Khan^25^ were ordered from Integrated DNA Technologies (Coralville, Iowa, USA). The reverse primer was used with HIV-RT enzyme for cDNA synthesis (RT step). Based on the primer binding location on the RNA template, we expected a cDNA product up to 1,181 nucleotides long. The forward and reverse primers were used in the amplification reaction (qPCR), with an expected amplicon size of 140 nucleotides. Primers and probe were resuspended in 1X Tris-EDTA Buffer (pH 8.0, Fisher Scientific; Waltham, Massachusetts, USA; BP2473100) to reach a concentration of 100 µM, then diluted with nuclease-free water to a 10 µM working concentration. 100 µM and 10 µM solutions were kept at -20°C, and the probe was protected from light by aluminum foil.

### PERT Assay Workflow

10 µL cDNA synthesis reactions contained 1X HIV-RT Buffer, 0.2 U/µL RNasin® Plus Ribonuclease Inhibitor (Promega; Madison, Wisconsin, USA; N2611), 0.2 mM Deoxynucleotide (dNTP) Solution Mix (New England Biolabs; Ipswich, Massachusetts, USA; N0447), 0.2 µM reverse primer, 83.4 nM MS2 RNA, nuclease-free water, and 1 µL HIV-RT (1 µL 1X RT Buffer for the “No HIV-RT” or NRT condition). Reactions were incubated at 42°C for 60 minutes followed by enzyme inactivation at 95°C for 5 minutes. After reaction completion, cDNA samples were diluted 2X in nuclease-free water before addition to the qPCR reaction using triplicates (three technical replicates) of each cDNA reaction. For all figures except where indicated (e.g., **Figure 4B** and **Figure 8A**), N represents technical qPCR replicates from split cDNA synthesis reactions (3 replicates from each individual, completed cDNA synthesis reaction).

For the amplification and detection step of the PERT assay, 25 µL qPCR reactions were prepared containing 0.025 U/µL One*Taq*® Hot Start DNA Polymerase (New England Biolabs; Ipswich, Massachusetts, USA; M0481), 1x One*Taq®* Standard Reaction Buffer (New England Biolabs; Ipswich, Massachusetts, USA; B9022), 0.2 mM dNTP Solution Mix, 0.4 µM forward primer, 0.2 µM reverse primer, 0.05 µM probe, nuclease-free water, and 5 µL of 2X diluted cDNA (5 µL nuclease-free water for the “No cDNA Template” or NTC condition). Amplification reactions consisted of 2 min at 95°C followed by 45 cycles of 15s at 95°C and 30s at 56°C. One set of negative controls (three replicates of NRT and NTC each) was included in each qPCR run; corresponding controls for the run are shown on each raw qPCR curve figure and standard curve. A detailed procedure is included in the Supplemental Material and results using this protocol can be found in **Figure 2, 3**, and **5**.

**Figure 1.**
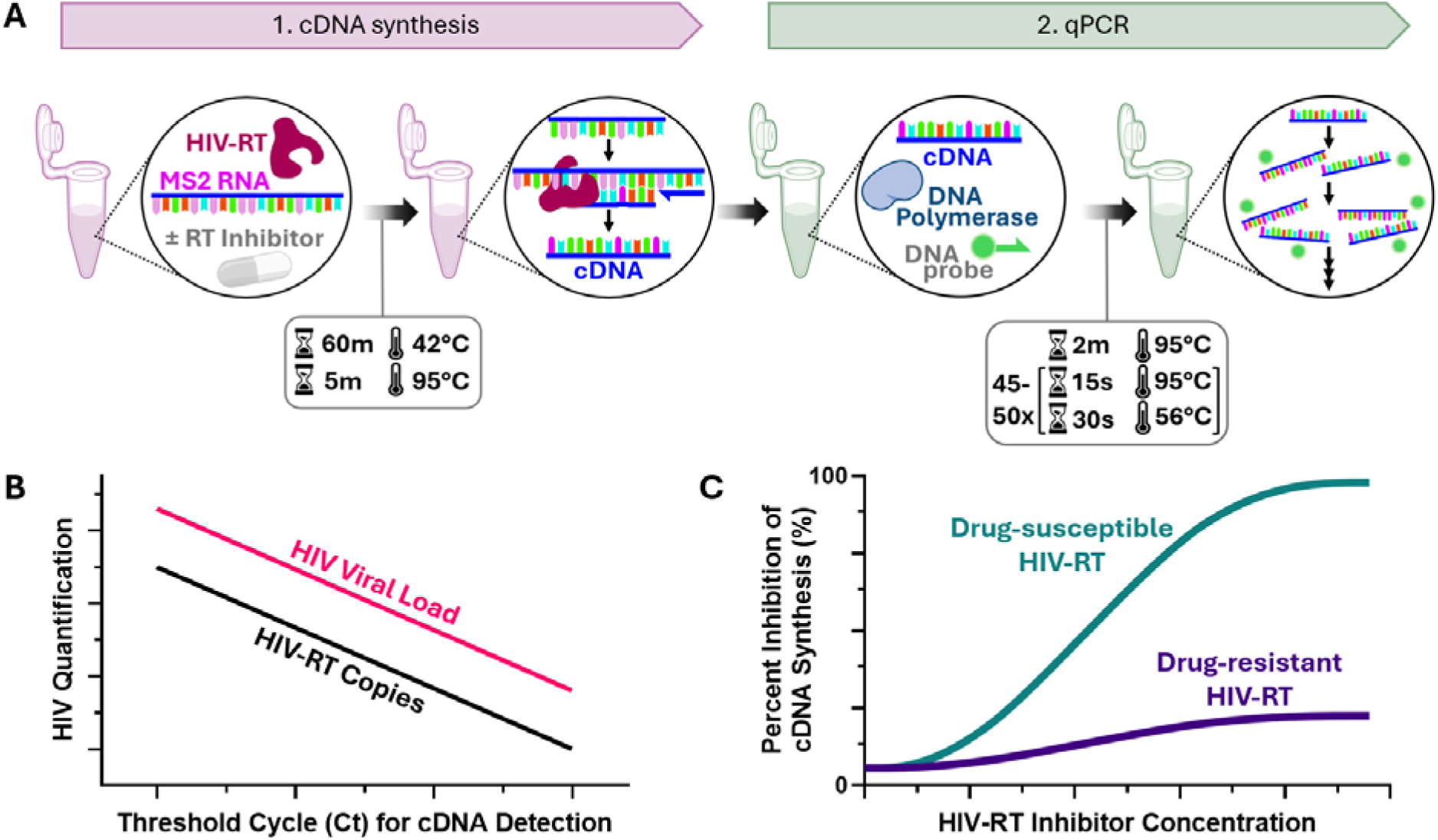
Overview and operating principle of PERT assay. **(A)** The first step in PERT is cDNA synthesis by HIV-Reverse Transcriptase (HIV-RT) enzyme in the presence and absence of drug, with the extent of synthesis dependent on the enzyme’s drug resistance/susceptibility. The second step is quantitative PCR (qPCR) for rapid and accurate measurement of the cDNA produced. **(B)** An example of standard curves generated using PERT. RT-qPCR provides linear quantification of HIV-RT, proportional to HIV viral load (∼50- 100 HIV-RT enzymes/virion and 2 HIV RNA molecules/virion), based on HIV-RT enzyme activity. **(C)** PERT distinguishes between drug-resistant and drug-susceptible HIV-RT enzymes based on the inhibition of cDNA synthesis in the presence of HIV-RT inhibitor drugs.

**Figure 2.**
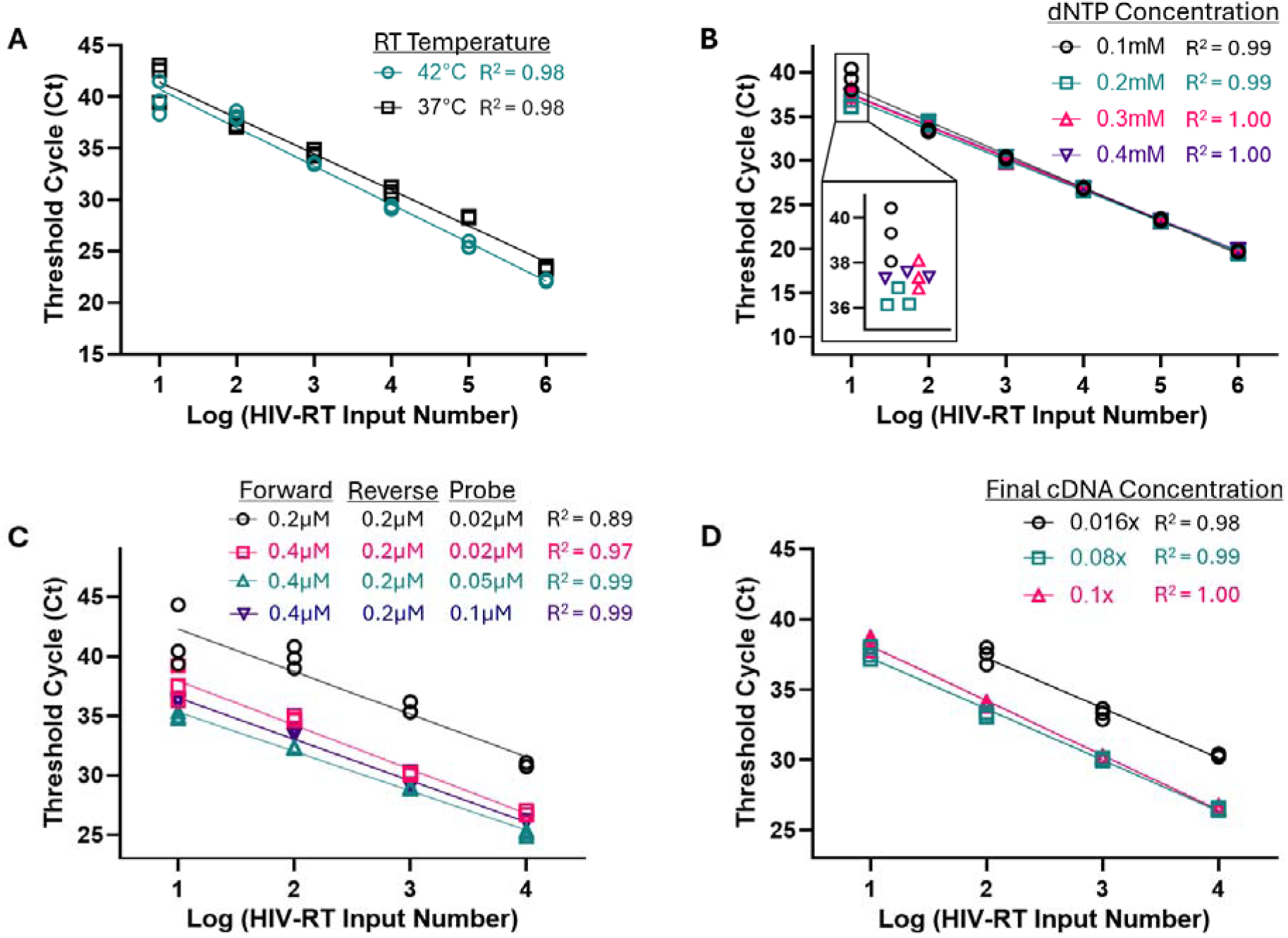
Optimization of the PERT assay for detection of low levels of HIV-RT enzyme. Standard curves for a range of input HIV-RT concentrations with the measured threshold cycle (Ct) for each condition: **(A)** RT reaction (cDNA synthesis) at 37°C and 42°C, **(B)** varied deoxynucleotide (dGTP, dATP, dCTP, and dTTP) concentration in the RT reaction, **(C)** qPCR primer and probe concentration, and **(D)** cDNA concentration in qPCR reactions. Optimal conditions for each experiment are shown in green. Experiments were completed by dividing each completed cDNA sample into three technical qPCR replicates (N=3).

**Figure 3.**
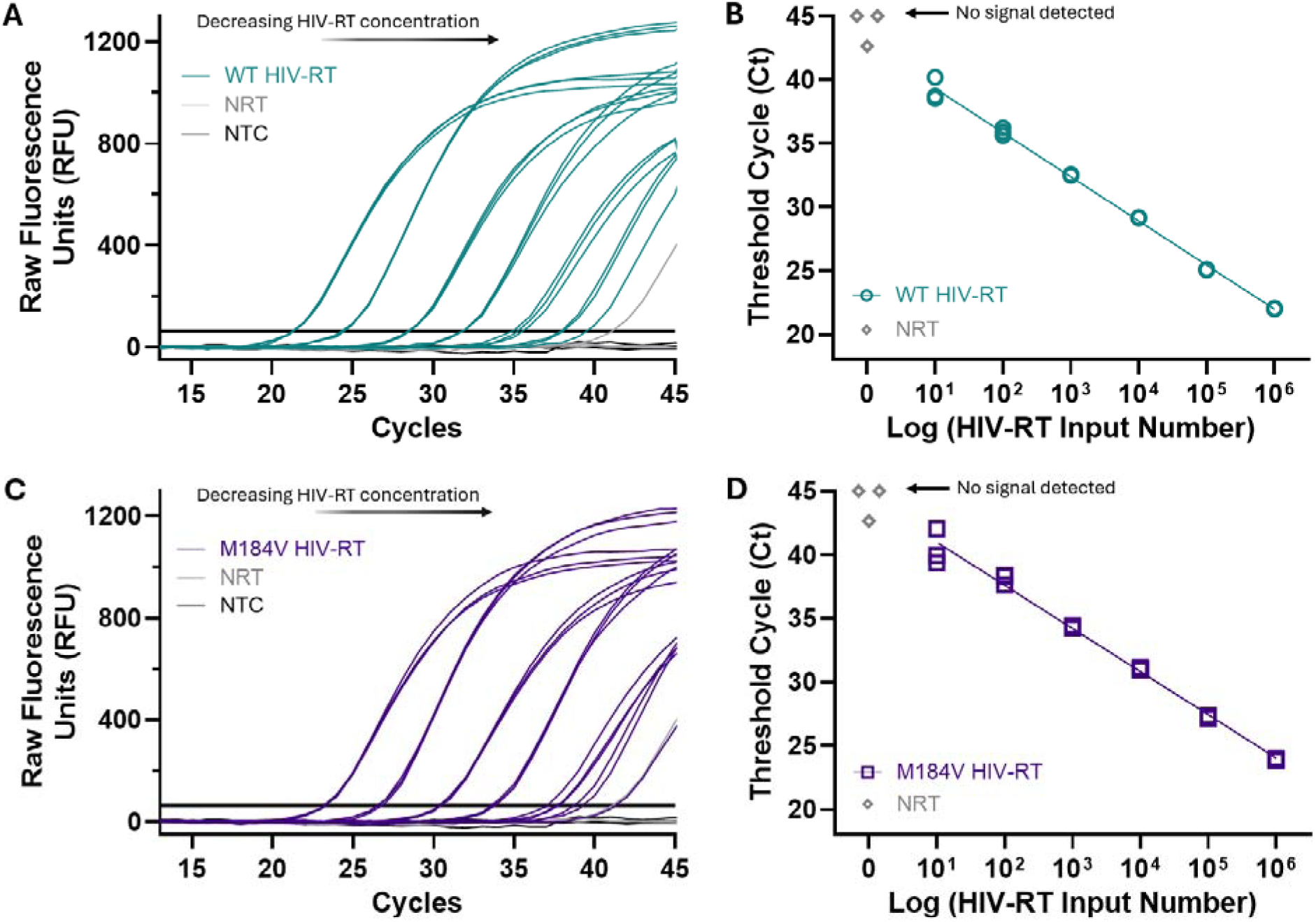
Sensitive detection of HIV-RT activity using an optimized PERT assay. (A,C) Raw qPCR curves for WT and M184V HIV-RT, respectively, and negative (NRT and NTC) controls, with Ct values increasing according to decreasing HIV-RT concentrations. **(B,D)** qPCR standard curves for WT and M184V HIV-RT showing linear quantification of HIV-RT enzyme and its activity within the range of concentrations tested (10 to 10^6^ molecules HIV- RT). Experiments were completed by dividing each completed cDNA sample into triplicates for qPCR (N=3), with WT and M184V samples and one set of negative controls run on the same qPCR plate.

**Figure 4.**
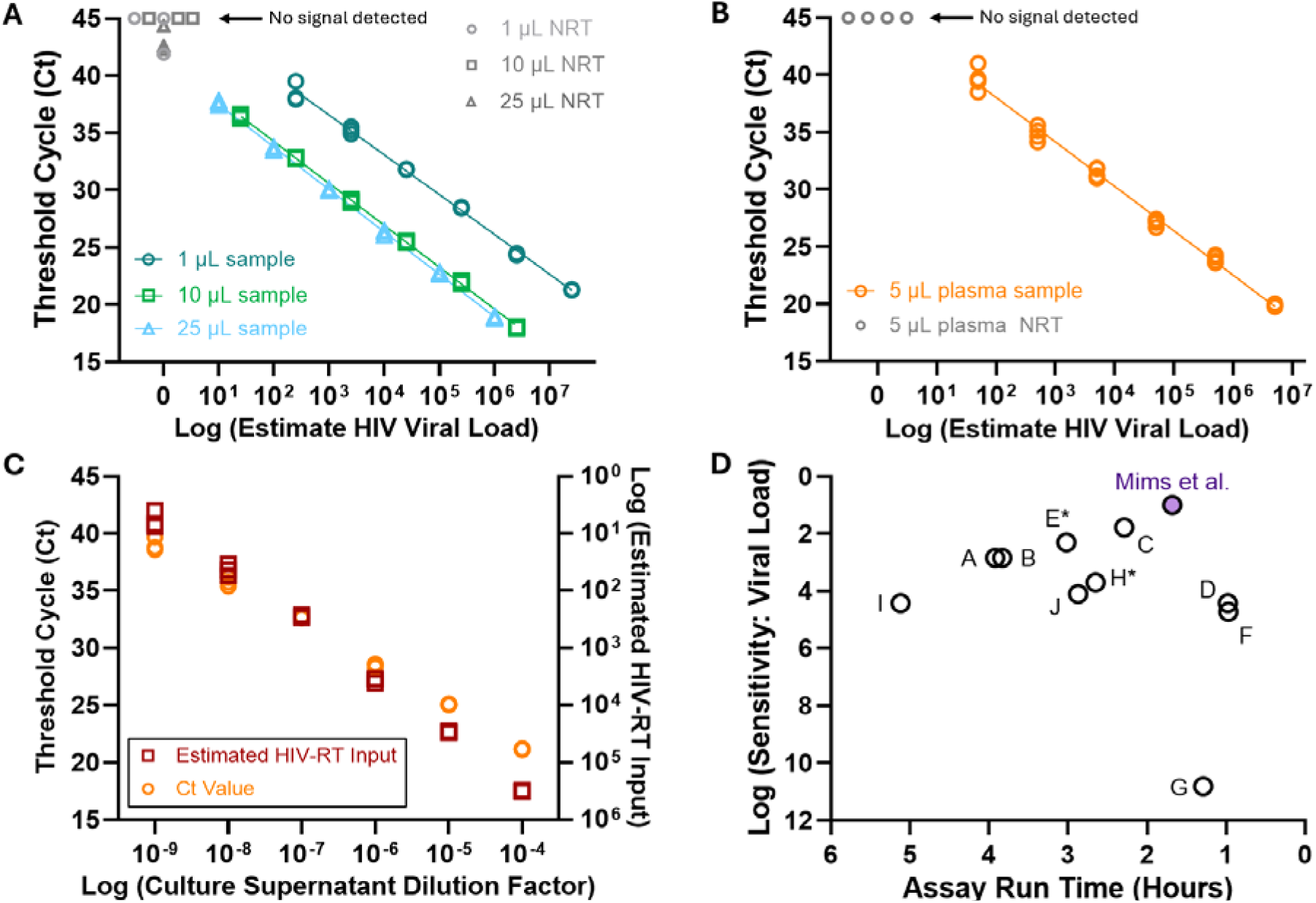
Detection of HIV-RT activity using larger sample volumes, plasma, and culture supernatant. **(A)** As input sample volume is increased from 1 to 10 or 25 µL HIV-RT, the PERT assay provides linear quantification of HIV-RT activity across the same range of concentrations (10 to 10^6^ HIV-RT), with increased volume providing increased sensitivity according to corresponding VL. **(B)** When spiking 1 µL HIV-RT into 5 µL plasma in the 20 µL cDNA synthesis reaction (using a modified RT Buffer), PERT detects down to 10 input HIV-RT with minimal amplification of negative controls. **(C)** Ct values and estimated HIV-RT input from each 5 µL HIV-1 cell culture supernatant sample (estimate HIV-RT input calculated from a standard curve of HIV-RT in buffer, **Supplementary** Figure 3C) show that the modified RT buffer can lyse HIV virions for direct testing of HIV-RT activity. Experiments were completed by dividing each completed cDNA sample into triplicates (**A, C**) or duplicates (**B**) for qPCR (N=3-4), with NRT and NTC controls included for each. **(D)** Comparison of our optimized PERT assay to previous demonstrations^22,25,33–40^ according to their run time and sensitivity in terms of VL detected. Full comparisons can be found in **Table S4**. *\*All VL levels were calculated using the assay’s reported HIV-RT sample volume for cDNA synthesis, assuming 80 HIV-RT/virion and 2 copies of RNA/virion (except Hoffman et al & Macchi et al, which were reported in terms of VL)*.

### Optimization of PERT for Measuring Low Concentrations of HIV-RT

For assay optimization, various conditions of the cDNA synthesis and qPCR reactions were modified, including dNTP concentration, cDNA synthesis reaction temperature, primer and probe concentrations, and dilution factor and volume of cDNA added to the qPCR reaction (**Figure 2, Supplementary Table 1**). All optimization experiments were performed using WT HIV-RT.

### PERT Assay with Increased Sample Volume

20 µL cDNA synthesis reactions (**Figures 4-8**) contained the same concentration of all components except 41.7 nM MS2 RNA and 10 µL HIV-RT (10 µL 1X HIV-RT Buffer for the NRT condition). After incubation for cDNA synthesis, cDNA reactions were not diluted before addition to the qPCR reaction, with triplicates of each cDNA reaction (5 µL cDNA for each replicate) used in the qPCR reaction. The probe concentration was increased to 0.1 µM for the qPCR reaction.

35 µL cDNA synthesis reactions (**Figure 4A**) contained the reduced RNA concentration as well as modified RT buffers: The 1X RT buffer used for diluting HIV-RT samples contained 50 mM Tris-HCl, 8 mM MgCl_2_, and 0.5% Triton X (no KCl or DTT) while the 1X RT buffer used for the cDNA synthesis master mix contained 50 mM Tris-HCl, 16.7 mM KCl, 5 mM DTT, and 1% Triton X (no MgCl_2_ and reduced KCl and Triton X). 25 µL HIV-RT samples were combined with 10 µL master mix for cDNA synthesis; 10 µL of each completed cDNA synthesis reaction was added to the qPCR reaction in triplicates (keeping the qPCR reaction volume at 25 µL total) and using the increased 0.1 µM probe concentration.

### PERT Assay with Plasma or Cell Culture Supernatant

The process listed in the **PERT Assay with Increased Sample Volume** for 20 µL cDNA synthesis reactions was utilized for testing spiked plasma and cell culture supernatant samples (**Figure 4B-C**), except reduced sample volumes in the 20 µL reaction. For plasma testing, 5 µL of plasma and 1 µL of HIV-RT were added to the reaction. After cDNA synthesis, cDNA samples were vortexed and centrifuged using a benchtop centrifuge to separate liquid cDNA from now- solid “gooped” plasma. 3 µL cDNA was used for each qPCR reaction; duplicates or triplicates of each cDNA sample were used. For HIV-1 cell culture supernatant, 5 µL of supernatant was used for sample input in the 20 µL cDNA synthesis reaction and 5 µL of each cDNA sample used in triplicate for qPCR. For plasma and culture supernatant, the modified buffer from the **HIV-RT Buffer for Direct Plasma and HIV-1 Culture Testing** section (containing increased Triton X and decreased DTT) was used for dilution of HIV-RT prior to mixing with plasma or dilution of culture supernatant as well as for the cDNA synthesis reaction master mix and NRT controls.

### Distinguishing Between Drug-Resistant and Drug-Susceptible HIV-RT

When distinguishing between resistant and susceptible HIV-RT with NNRTI (**Figure 7D**), no changes were made to the reagent concentrations other than reduction of water volume to account for the addition of the RT inhibitor DOR. For distinguishing between resistant and susceptible HIV-RT with nucleotide analog drugs (NRTIs and NRTTI), the processes listed above in the **PERT Assay Workflow** section or **PERT Assay with Increased Sample Volume** section were followed except for the changes described below. The concentration of nucleotide for which the drug is an analog was decreased (keeping all other dNTPs at 200µM) (New England Biolabs; Ipswich, Massachusetts, USA; N0446S) to enable greater detection of inhibition by the nucleotide analog drugs. Specifically, 50 µM dCTP was used in the reactions with the C analog drug (3TC-TP: **Figure 5-6**) and 50 µM dATP used in the reactions with A analog drugs (ISL-TP and TFV-DP: **Figure 7A-C**). Additionally, before adding HIV-RT to the cDNA Reaction Mix, 1-2 µL of RT inhibitor was added to the reaction (1-2 µL of water for no-drug conditions) for 10-20 µL cDNA synthesis reactions.

**Figure 5.**
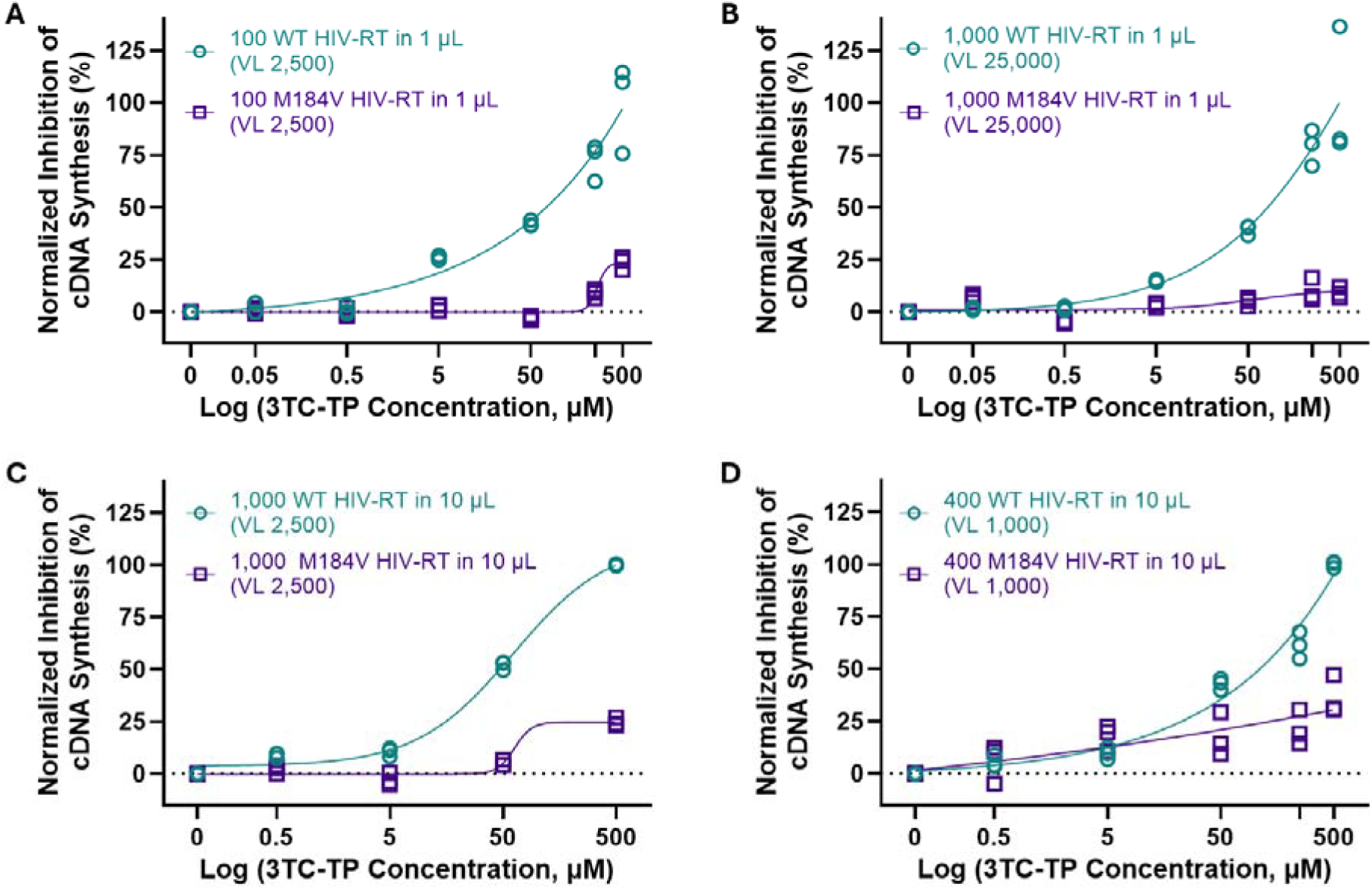
PERT distinguishes between drug-resistant and drug-susceptible HIV-RT in the presence of 3TC-TP. Normalized inhibition curves for the PERT assay comparing WT and M184V HIV-RT enzyme as 3TC-TP concentration increases from 50 or 500 nM to 500 µM in 10-fold increments. Analysis of normalized inhibition of cDNA synthesis shows that the assay can distinguish between 100 molecules of WT and 100 molecules of M184V HIV-RT using 1 µL HIV-RT sample **(A)**, between 1,000 WT and 1,000 M184V HIV-RT a 1 µL HIV-RT sample **(B)** or 10 µL sample **(C)**, and between 400 WT and 400 M184V HIV-RT in a 10 µL HIV-RT sample **(D)**. Experiments were completed by dividing each completed cDNA sample into triplicates for qPCR (N=3), and nonlinear regression was performed for each curve.

**Figure 6.**
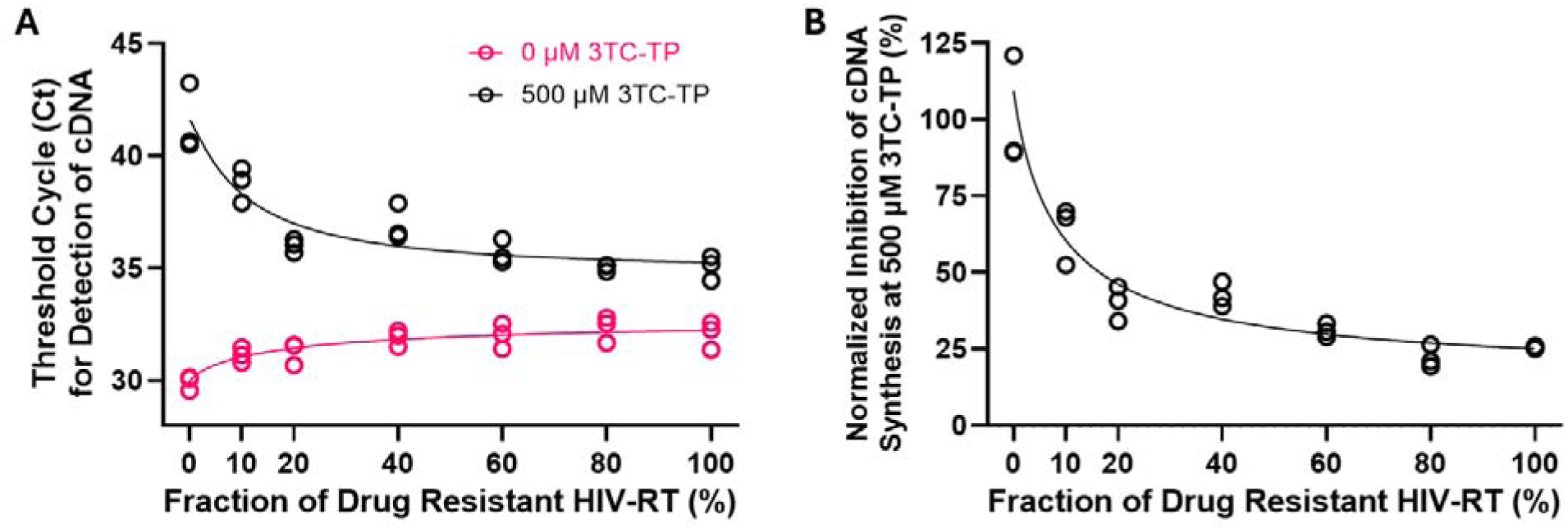
Quantitative detection of NRTI drug resistance by PERT using heterogeneous HIV-RT fractions. **(A)** Ct values for heterogeneous mixtures of WT and M184V HIV-RT at 0 µM 3TC-TP show small losses in activity with increased presence of M184V HIV-RT, while Ct values at 500 µM 3TC-TP show less reduced activity with increased presence of M184V HIV- RT. **(B)** Normalized inhibition values for each drug-resistant fraction at 500 µM 3TC-TP show that the assay can distinguish between WT and mutant fractions as low as 10%, confirmed using one-way ANOVA (p < 0.0001) and post-hoc multiple comparisons test (**Table S6**). Experiments were completed by dividing each completed cDNA sample into triplicates for qPCR (N=3), and nonlinear regression was performed for each curve fit.

**Figure 7.**
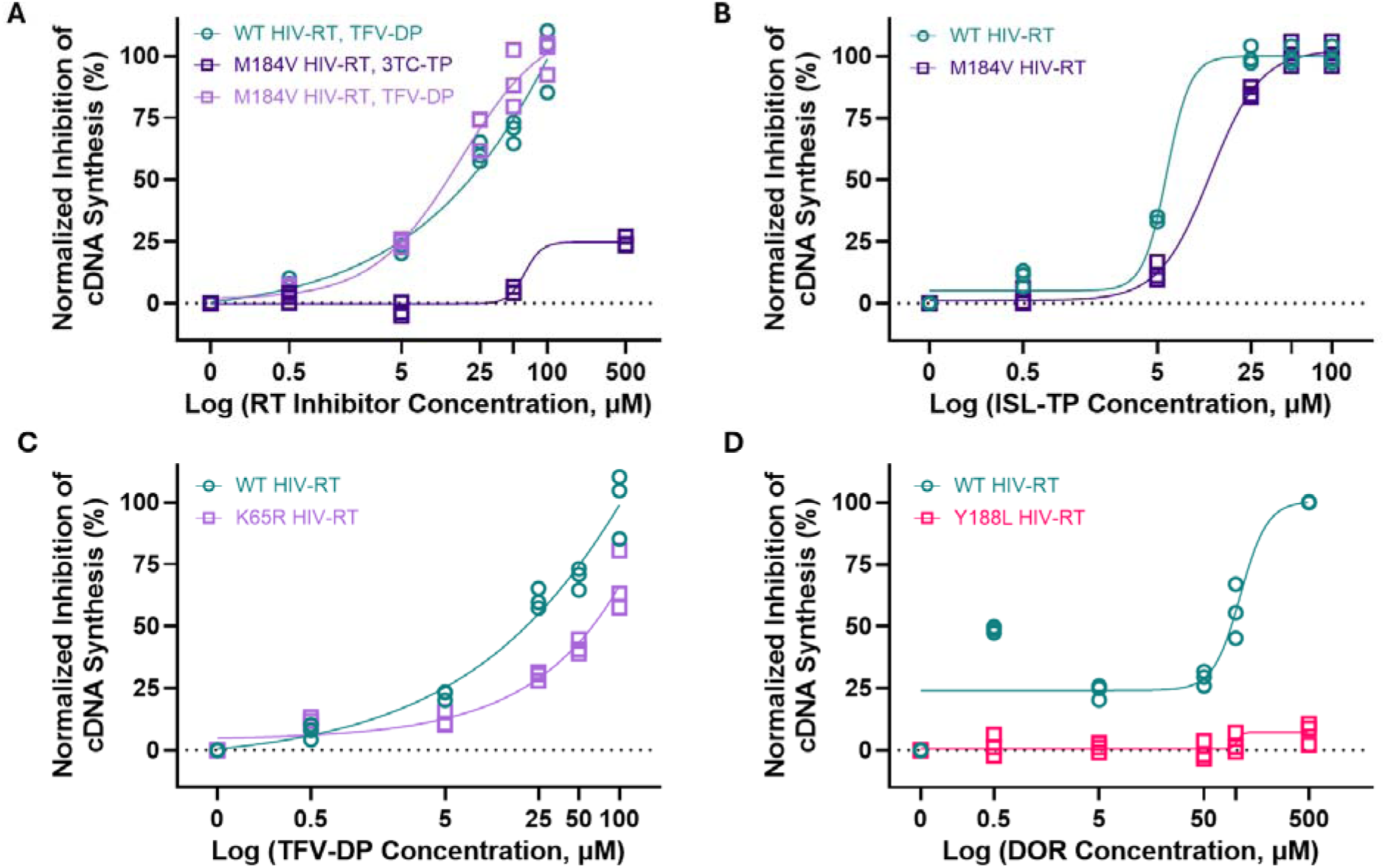
PERT provides specific distinction between drug-resistant and drug- susceptible HIV-RT. **(A)** Normalized inhibition curves for the PERT assay comparing WT and M184V HIV-RT enzyme in the presence of TFV-DP show that the assay can specifically distinguish between resistant and susceptible HIV-RT mutations. **(B-D)** Analysis of normalized inhibition of cDNA synthesis for the M184V, K65R, and Y188L HIV-RT mutations show distinction between WT and resistant mutations using ISL-TP, TFV-DP, and DOR, respectively (**Table S7**). Experiments were completed using 1,000 input HIV-RT in a 10 µL sample (corresponding to a VL ∼2,500 RNA/mL) and by dividing each completed cDNA sample into triplicates for qPCR (N=3). Nonlinear regression was performed for each curve.

**Figure 8.**
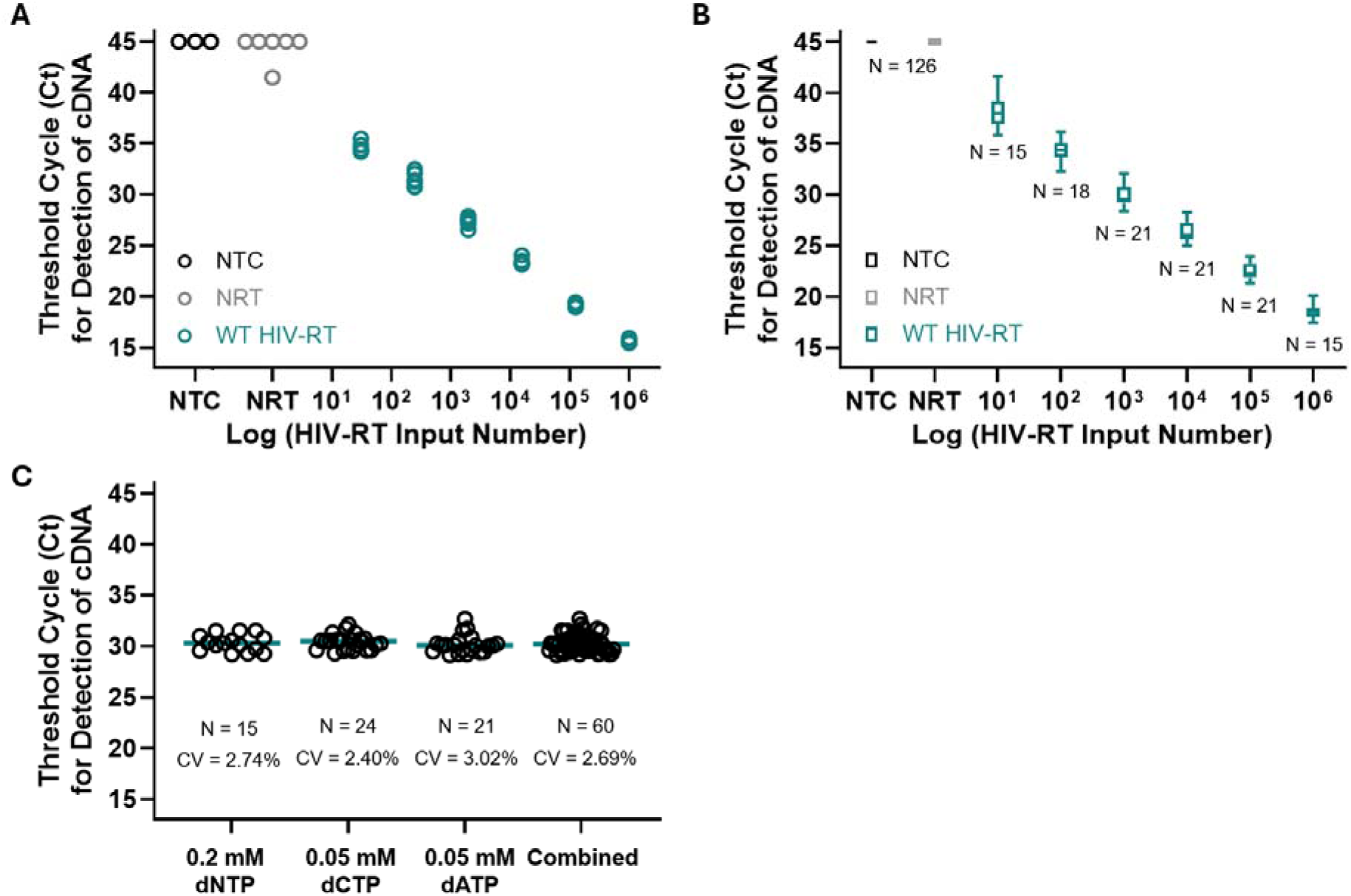
PERT provides highly reproducible quantification of HIV-RT activity for VL measurement and DRT. **(A)** Intra-assay variation analysis using 3 cDNA replicates (two qPCR replicates for each, giving N=6 for each HIV-RT concentration) shows that cDNA replicates do not introduce variability (< 3% coefficient of variation, or CV, for all HIV-RT samples) with 1 NRT control detected. **(B)** Inter-assay variation—combining results from > 6 months—shows that the PERT assay tolerates changes in sample volume, reagent stocks, machines, and seasonal changes, with < 5% CV for all samples, including NRT. **(C)** Inter- assay variation for detection of 1,000 HIV-RT at varied dNTP concentration (equal dNTP or reduced amount of the specific dNTP listed) shows that using the decreasing nucleotide concentration for analog drugs does not increase assay variability (∼ 3% CV). See **Table S8** for a full analysis of the intra- and inter-assay variation.

### Detecting Drug Resistant HIV-RT Fraction in Heterogeneous Mixtures

WT and M184V HIV-RT enzyme were mixed to obtain 10, 20, 40, 60, and 80% resistant fractions. 0% and 100% resistant fractions were represented by WT-only and M184V-only fractions, respectively. As in the prior section, the dCTP concentration was decreased in the cDNA synthesis reaction; 50 µM dCTP was used in the RT reactions, both with and without 3TC-TP (**Figure 6**). 10 µL of each HIV-RT enzyme mixture and 2 µL of 3TC-TP were added to each 20 µL cDNA synthesis reaction.

### Equipment, Statistical Analysis, and Data Representation

cDNA synthesis was performed on a heat block (Benchmark Scientific BSH1002 Digital Dry Bath Heater) or thermal cycler (Bio Rad CFX96 Touch Real-Time PCR Detection System); enzyme inactivation following cDNA synthesis as well as qPCR reactions were performed on the thermal cycler (CFX96). Raw amplification curves as well as threshold cycle (Ct) values were obtained from the thermal cycler, with analysis and calculations performed using Microsoft® Excel. The auto-generated threshold was utilized for each optimization experiment (**Figure 2** and **Supplementary Figure 5**) and the baseline threshold set to 100 for all other experiments and figures. All statistical evaluations and graphical representations were performed using GraphPad Prism version 10.6.0 for Windows (GraphPad Software Inc.).

To create standard curves for experiments with varied HIV-RT concentrations, we plotted log10(HIV-RT molecules) against the Ct at which the cDNA product was detected by qPCR and performed a linear regression. When analyzing enzyme inhibition, we calculated ΔCt values as the difference between drug and no-drug conditions for each sample and normalized this value to each assay’s maximum inhibition condition as follows:

Normalized Inhibition of Sample %

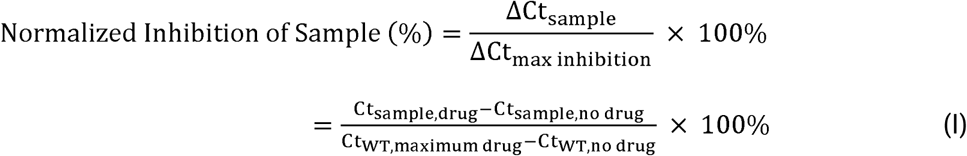

We fit four-parameter logistic regression curves for the Ct, ΔCt, and normalized inhibition plots (all default options). We determined statistically significant differentiation of inhibition of sample compared to the WT condition using one-way ANOVA followed by a post-hoc Bonferroni’s or Dunnett’s multiple comparisons test, for homogenous or heterogeneous enzyme fractions, respectively.

Any cDNA sample replicates—including for NRT controls—that were not detected within the qPCR cycles were set to a value matching the assay’s maximum number of cycles (45 cycles for all figures except for 50 in **Figure S5**).

## RESULTS AND DISCUSSION

### Overview and Operating Principle of PERT

PERT assays can measure both HIV VL and resistance to RT inhibitors based on the activity of HIV-RT enzyme (**Figure 1**). The first step in PERT is an *in vitro* reverse transcription reaction during which HIV-RT synthesizes varying amounts of cDNA depending on HIV-RT enzyme concentration and resistance/susceptibility to reverse transcriptase inhibitors. The second step is the amplification of synthesized cDNA by qPCR resulting in a fluorescence readout. Using this two-step RT-qPCR format, PERT can quantify HIV VL as a function of the cDNA synthesis activity of HIV-RT (∼50-100 HIV-RT enzymes per virion^25–29^ and 2 HIV RNA copies per virion) in the no-drug condition (**Figure 1B**). Larger HIV-RT input samples will produce greater cDNA quantities for amplification by qPCR and lower cycle threshold (Ct) values. When PERT is performed in the presence of drugs that target HIV-RT, drug-susceptible HIV-RT enzymes are strongly inhibited in their cDNA synthesis activity, while drug-resistant HIV-RT enzymes show weak or delayed inhibition of cDNA synthesis (**Figure 1C**).

### Optimization of PERT for Measuring Low Concentrations of HIV-RT

We optimized PERT to detect very low levels of WT HIV-RT enzyme (**Figure 2**). We aimed to achieve Ct values <45 for all conditions containing HIV-RT enzyme, without detecting “No HIV-RT” (NRT) or “No cDNA Template” (NTC) controls. Linear regression was performed for each dataset and the coefficient of determination (R^2^) reported; linear regression equations and calculated efficiencies for each are reported in **Table S1** and raw qPCR curves are reported in **Supplementary Figure 1**.

We first examined the impact of the RT reaction temperature on Ct values since HIV-RT is active between ∼25–45°C (**Figure 2A**). We chose 37°C and 42°C as our test temperatures since they are closer to the optimal operating conditions for the enzyme and, in future testing in hot climates, would be less likely to require active cooling. cDNA synthesis at 42°C produced slightly lower Ct values compared to 37°C; however, there were minimal differences between the two temperatures.

We also examined the impact of deoxynucleotide (dNTP) concentration on detection of HIV-RT activity (**Figure 2B**). Starting with 83.4 nM of MS2 RNA template in the RT reaction, we systematically varied dNTP concentration to provide ∼30 to 150-fold excess dNTP per template to synthesize full-length amplicons starting from the reverse primer. Across the range of dNTP concentrations screened, the only impact we observed on Ct values was when detecting 10 molecules of HIV-RT. Since our goal was to detect drug resistance at the lowest HIV-RT copy number and lowest dNTP concentration achievable—to strike a balance between non-limiting conditions and dNTP-RT inhibitor competition—we selected 0.2 mM of each dNTP moving forward with PERT. Calculations for determining the ratio of each dNTP to template are shown in **Table S2**.

Next, we optimized qPCR conditions by modifying primer and probe concentrations (**Figure 2C**). We used the same primers and probe in the qPCR assay design by Ma and Khan^25^ that had the probe binding to the DNA template only a few nucleotides past the forward primer.

While this design could potentially provide higher analytical sensitivity due to the increased probability of probe binding prior to extension of the DNA strand by DNA polymerase, there is also a chance of the probe interfering with and destabilizing forward primer binding.^30^ Therefore, we kept forward primer concentration greater than probe and reverse primer concentrations, to mitigate any losses in sensitivity from probe binding. Increasing forward primer to 0.4 µM while keeping reverse primer at 0.2 µM showed drastic improvements in Ct values. Meanwhile, increasing the probe concentration from 0.02 µM to 0.05 µM improved Ct values (by an average of 2.01 ± 0.65 cycles), but performance approached baseline conditions again at 0.1 µM probe concentration. Therefore, we chose 0.4 µM, 0.2 µM, and 0.05 µM as the optimal concentrations for forward primer, reverse primer, and probe. Using these conditions, we could run 45-50 cycles of PCR amplification with minimal non-specific amplification/signal from negative (NRT and NTC) controls.

Because prior PERT assays were sensitive to the final cDNA volume/concentration in the qPCR reaction,^21^ we also examined the impact of dilution and volume of cDNA product from the RT reaction prior to the qPCR step on the overall assay performance (**Figure 2D**). 10 µL cDNA synthesis reactions were either not diluted, diluted 2X, or diluted 5X (with nuclease-free water) prior to addition to the amplification reaction for final cDNA concentrations in the qPCR assays of 0.08X, 0.1X, and 0.016X, respectively. For the undiluted and 5x dilution, we added 2 µL of cDNA product to 23 µL of qPCR master mix; for the 2x dilution, we added 5 µL of cDNA product to the 20 µL of qPCR master mix. We obtained nearly identical standard curves at cDNA concentrations of 0.08X and 0.1X, suggesting that this slight difference did not significantly impact assay conditions. Our results show that this PERT assay is flexible in the sample volume requirements and could support further modifications.

Using the optimized assay conditions described above, we tested a range of WT and M184V HIV-RT concentrations (**Figure 3**). When spiking ≥ 10 molecules of HIV-RT into the cDNA synthesis reaction, we detected cDNA products within 45 cycles of qPCR. All replicates containing HIV-RT enzyme were detected; NRT replicates would occasionally be detectable (e.g., **Figure 3**), but no more than 2 of 3 replicates ever amplified and no NTC replicate was ever detected (**Figure 8**). The Ct values for M184V HIV-RT were slightly higher than those of WT HIV-RT, as expected, due to the reduced viral fitness of mutant HIV-RT.^31,32^ Viral fitness— defined as the ability to adapt and reproduce in a defined environment and shown to correlate with reduced natural nucleotide incorporation efficiency—therefore only slightly affected the cDNA synthesis ability of the mutant HIV-RT compared to WT. The dynamic range of the optimized PERT assay extends over at least 6 log_10_ (**Figure 3B, 3D**), with linear quantification of HIV-RT enzyme and its activity within the range of concentrations tested (10 to 10^6^ molecules HIV-RT). The linear regression equations, coefficients of determination, and efficiency of each reaction (calculated from the slope of the equation and reported with 95% confidence intervals) for WT and M184V HIV-RT were Y 3.469x % 42.8 (R^2^ = 1.0, efficiency = 94.21% [90.01, 98.90]) and (R^2^ = 0.99, efficiency = 97.24% [90.39, 105.1]), respectively, and are reported in **Table S1**.

### Increasing HIV-RT Sample Volume Improves PERT Assay Sensitivity

Although the optimized PERT assay presented here detects 10 HIV-RT input molecules, this assay iteration used 1 µL enzyme sample in a 10 µL reaction, which limits how much RT enzyme is available to detect and corresponds to a VL ∼250 copies/mL. We hypothesized that the reaction volume could be scaled up, enabling use of a 10-25 µL sample volume and providing HIV-RT detection corresponding to a VL of ∼25 and 10 copies/mL, respectively. See **Table S3** for the corresponding VL according to various HIV-RT quantities and sample volumes. We tested this adapted assay format by spiking ≥ 10 WT HIV-RT in a 10 or 25 µL sample for input into the 20 or 35 µL cDNA synthesis reaction and still detected all replicates containing HIV-RT enzyme within 45 cycles of qPCR (**Figure 4A**), again with only occasional NRT replicates detectable. Therefore, with the modified format using increased sample volume in the cDNA synthesis reaction, this PERT assay showed detection of HIV-RT down to a corresponding VL of ∼25 or 10 copies RNA/mL. Raw qPCR curves as well as standard curves according to HIV-RT input quantity for the varied sample volumes can be found in **Supplementary Figures 2** and **3**, respectively. Linear fit equations, R^2^ values, and efficiencies for each can be found in **Table S1**.

### Extraction-free Plasma and Cell Culture Supernatant Testing with PERT

Prior work by Gerardo García-Lerma and Walid Heneine suggests that addition of detergent to plasma to lyse virions followed by direct addition to a phenotypic enzymatic assay could provide extraction-free detection of HIV-RT activity.^22^ To demonstrate proof-of-concept testing of extraction-free HIV-RT activity detection in our hands, we spiked 10 to 10^6^ HIV-RT into HIV-negative plasma and utilized the 20 µL cDNA synthesis reaction to examine inhibition of the RT-qPCR assay by the presence of plasma (**Figure 4B**). We still observed linear detection of HIV-RT activity down to 10 input HIV-RT, with minimal detection of NRT samples (5 µL plasma and 1 µL 1X RT Buffer).

When testing assay performance with plasma, we noticed that with the original buffer concentrations, the plasma-containing samples were nearly completely solid “goop” after cDNA synthesis (**Supplementary Figure 4C**), with samples un-pipetteable for transfer of produced cDNA to qPCR reactions. With these solid samples, we added water to the goop and vortexed them to release cDNA then transferred the liquid to qPCR, but assay performance was poor and limited by NRT amplification (**Supplementary Figure 4B-C**). After decreasing the DTT concentration (from 5 mM to 2.5 mM in the 1X RT Buffer) and increasing the Triton X concentration (from 0.16% to 1.6% in the 1X RT Buffer), we could detect 1 µL of HIV-RT spiked into 5 µL plasma and added directly to a 14 µL cDNA master mix (**Figure 4C** and **Supplementary Figure A,C**). The buffer modifications allowed increased separation of liquid cDNA sample from solid plasma goop following cDNA synthesis while minimizing assay inhibition. With this adapted buffer, we could obtain sufficient volumes of un-gooped liquid to perform duplicate or triplicate 3 µL synthesized cDNA samples for downstream addition to qPCR.

The modified RT Buffer used for plasma testing also enabled direct lysis of HIV virions: We diluted NL4-3 HIV-1 cell culture supernatant in the modified 1X RT Buffer and added 5 µL of diluted supernatant to the 20 µL cDNA synthesis reaction (followed by triplicates of 5 µL cDNA samples added to the qPCR reaction) for detection of HIV-RT activity (**Figure 4C**). Using a standard curve from HIV-RT spiked buffer (**Supplementary Figure 3C**), we calculated the estimated HIV-RT input for each culture supernatant sample. While this shows proof-of-concept ability to directly lyse virions for detection of HIV-RT activity by PERT, additional validation with clinical samples is required for both plasma and culture samples.

Raw qPCR curves as well as standard curves according to HIV-RT input quantity for the plasma and culture supernatant samples can be found in **Supplementary Figures 2** and **3**, respectively. The linear fit equation, R^2^ value, and efficiency for plasma testing can be found in **Table S1**.

### Comparison with Prior PERT assays

We compared our optimized assay to prior work focused on detecting HIV-RT activity,^22,25,33–40^ in terms of the VL that corresponds to the molecules of HIV-RT detected per reaction (**Figure 4D**). This comparison highlights the tradeoff between assay sensitivity and turnaround time. Our PERT assay was more sensitive than prior assays in measuring molecules of HIV-RT or VL while also providing one of the shortest assay times, with the difference primarily due to the length of the cDNA synthesis (RT) step. A full comparison of specimen type tested, readout method, sensitivity, and assay run time as well as the reference for each letter in **Figure 4D** can be found in **Table S4**.

Although the classic approaches for measuring VL focus on HIV RNA, measuring HIV-RT by its activity instead provides the benefits of increased sensitivity with 50-100 molecules of HIV-RT compared to 2 copies of RNA per virion, conserved activity among subtypes, and increased accuracy by only quantifying virally competent HIV.^17,19,20,41^ Due to the wide range of HIV-RT molecules (50-100) that can be found within each HIV virion, this quantification of HIV-RT is an estimate of VL. However, prior studies have shown strong correlation between Ct values from qPCR-based HIV-RT enzyme activity assays and plasma VL.^17,41^ VL tests are expected to quantify down to 200 copies/mL (although ideally down to undetectable levels, i.e., 20-50 copies/ mL).^3,42^ Detecting 10 molecules of HIV-RT in a 10 µL PERT reaction corresponds to ∼20-40 copies HIV RNA/mL (assuming 50-100 HIV RT enzymes per virion^25–29)^ and the presence of virus only in plasma (∼50% of blood), suggesting that this optimized PERT assay may be able to achieve the required sensitivity levels for VL monitoring alongside clinical resistance testing.

### Distinguishing between Drug-Resistant and Drug-Susceptible HIV-RT

Inclusion of RT inhibitor in the RT synthesis step inhibits HIV-RT activity,^43^ reducing cDNA synthesis activity by the enzyme, but drug-resistant HIV-RT will be less inhibited (compared to drug-susceptible HIV-RT). Therefore, by examining the ability of an HIV-RT sample to synthesize cDNA in the presence of an RT inhibitor, we can distinguish between resistant and susceptible HIV-RT. We first tested the well-characterized M184V mutation and the first-line NRTI 3TC-TP. Since 3TC-TP is a dCTP analog that competes with the nucleotide for incorporation during cDNA synthesis, we varied dCTP concentration during the RT step to increase the distinction between drug and no-drug conditions^44^ (**Supplementary Figure S5**). We kept all other dNTP (dGTP, dATP, and dTTP) concentrations at 200 µM and decreased dCTP concentration to 100, 50, and 5 µM and investigated changes to baseline (no drug) Ct values and inhibition of WT HIV-RT by 500 µM 3TC-TP. Lower dCTP concentrations resulted in increased inhibition of HIV-RT since there was more opportunity for 3TC-TP incorporation. The differences in Ct between the drug and no-drug concentrations (ΔCt) were 17.8, 11.1, 11.2, and 9.1 for the 5, 50, 100, and 200 µM dCTP conditions, respectively. These results illustrate the greater inhibition of cDNA synthesis by 3TC-TP when there is less competition from dCTP. However, baseline HIV-RT activity was slightly inhibited at dCTP concentrations ≤ 50 µM. Since full-length cDNA may be up to 1,181 nucleotides long, 5 µM dCTP concentrations likely resulted in limiting reagent conditions (**Table S2**). Consequently, although 5 µM dCTP provided the largest ΔCt between drug and no-drug conditions, the number of qPCR cycles had to be extended past 45 for detection of reactions containing 3TC-TP. Even with extension to 50 cycles, those Ct values were near the assay’s limit of detection, adding potential risk of not detecting mutant HIV-RT samples with low levels of activity. Therefore, we operated at a dCTP concentration of 50 µM (or 50 µM dATP for dATP analog drugs) when running experiments to distinguish between drug and no-drug conditions while reducing limiting reagent effects.

We used the PERT assay to distinguish between resistant (M184V) and susceptible (WT) HIV-RT at a range of 3TC-TP concentrations (**Figure 5**, with raw qPCR curves and plots of Ct values in **Supplementary Figure 6** and **7**). We initially demonstrated this distinction at 100 and 1,000 WT or M184V HIV-RT using the original assay format of 1 µL HIV-RT sample in a 10 µL cDNA synthesis reaction (**Figure 5A** and **5B**). We quantified the inhibition of cDNA by calculating ΔCt between the no-drug control and each drug concentration normalized to the assay’s maximum inhibition condition of the WT HIV-RT in the presence of 500 µM 3TC-TP (Equation I). WT HIV-RT showed minimal inhibition at low drug concentrations and significant inhibition at high drug concentrations. In contrast, M184V HIV-RT showed minimal inhibition across drug concentrations. We performed a one-way ANOVA (p<0.0001) then compared each M184V drug condition to the respective WT drug condition using a post-hoc Bonferroni’s multiple comparison test and showed distinction (by normalized inhibition) between resistance and susceptible HIV-RT at 3TC-TP concentrations ≥ 5 and 50 µM for 100 and 1,000 HIV-RT, respectively (**Table S5**). At this sample volume, 100 and 1,000 input HIV-RT correspond to a VL of ∼2,500 and ∼25,000 copies RNA/mL, respectively (**Table S3**).

To increase the sensitivity in terms of corresponding VL, we also tested distinction of WT and M184V HIV-RT at the larger volume assay format (10 µL HIV-RT sample in a 20 µL cDNA synthesis reaction). At this increased sample volume, 1,000 input HIV-RT corresponds to a VL of ∼2,500 copies RNA/mL and 400 HIV-RT to a VL of ∼1,000 copies RNA/mL (**Table S3**), which puts us within the sensitivity requirement range of the WHO TPP (1,000-5,000 copies RNA/mL). Although we observed greater variation in the data at 400 input HIV-RT, likely because we were operating closer to the assay’s limit of detection, we could still statistically distinguish between WT and M184V at concentrations of 3TC-TP ≥ 0.5 and 50 µM for the 1,000 and 400 input HIV-RT, respectively (p<0.0001 by one-way ANOVA, **Figure 5C** and **5D, Table S5**). **Supplementary Figure 8** contains a comparison of the difference in Ct (ΔCt) and difference in normalized inhibition for each HIV-RT input quantity, sample volume, and corresponding VL. **Supplementary Figures 6** and **7** show the raw qPCR curves and plots of Ct values, respectively, for WT and M184V.

### Detecting Drug Resistant HIV-RT Fractions in Heterogenous Mixtures

Given the relationship between VL and HIV-RT activity,^17,41^ we assumed that a similar 20% threshold from sequencing methods (RNA-based) would apply to phenotypic DRT, and we examined the ability of the PERT assay to achieve this by combining WT and M184V HIV-RT in varied ratios (1,000 total molecules of HIV-RT in a 10 µL HIV-RT sample, corresponding to a VL of ∼2,500 HIV RNA/mL) and measuring the cDNA synthesis ability of each ratio in the presence of 3TC-TP (**Figure 6**). See **Figure S9** for raw qPCR curves of the drug resistance fractions.

At the baseline (without any drug), Ct values slowly increased as the fraction of mutant HIV-RT increased (**Figure 6A**), as expected with decreases in viral fitness and cDNA synthesis of M184V HIV-RT. At 500 µM 3TC-TP, there was a markedly different trend, with the WT (0% resistant fraction) samples having very high Ct values, representing significant inhibition, while the mutant (100% resistant fraction) samples showed minimal inhibition (matching the results seen in **Figure 5** for 100% WT and 100% M184V HIV-RT samples). To demonstrate that a single 3TC-TP concentration can be used to distinguish between WT and mutant HIV-RT (at clinically relevant mixtures with WT), we plotted the normalized inhibition values at the 500 µM drug condition according to resistant fraction percentage i.e., setting the ΔCt between 0 and 500 µM 3TC-TP in the WT (0% mutant fraction) as 100% inhibition in Equation I (**Figure 6B).** This drug concentration was chosen because it provided the largest distinction by difference in inhibition between WT and mutant samples (see **Figure 5**) and therefore the greatest window for distinction of resistant enzyme fractions. At 0%, 10%, 20%, 40%, 60%, 80%, and 100% mutant fraction, the normalized inhibition values of cDNA synthesis were 100 ± 18.2, 63.5 ± 9.6, 40.1 ± 5.7, 42.6 ± 4.1, 31.0 ± 2.2, 22.3 ± 3.7, and 25.5 ± 0.46, respectively. The normalized inhibition values at each of the mutant fractions tested were statistically distinguishable from WT (0% resistant HIV-RT) using a one-way ANOVA and post-hoc multiple comparisons test (**Table S6**).

### Drug Resistant Testing with Other HIV-RT Mutations and RT Inhibitors

Although the M184V mutation’s resistance to 3TC-TP^45^ is particularly well-characterized and therefore a good starting point for optimization and proof-of-concept DRT, many additional HIV-RT mutations confer resistance to other RT inhibitors, across different classes of inhibitors. Therefore, to assess the ability of the PERT assay to specifically detect resistance, we first tested the cDNA synthesis activity of the M184V HIV-RT in the presence of TFV-DP, to which the M184V mutation confers no resistance (and slight increases in susceptibility^46^). We compared inhibition of M184V with 3TC-TP and TFV-DP using 1,000 input HIV-RT in a 10 µL HIV-RT sample (**Figure 7A**). M184V HIV-RT showed slight increases in susceptibility to TFV-DP compared to WT; when using post-hoc multiple comparisons test, WT and M184V were only distinguishable at 50 µM TFV-DP (**Table S7**).

In addition to 3TC-TP resistance, the M184V mutation also confers low-level resistance (4-5- fold loss in susceptibility) to the newly approved NRTTI Islatravir.^47,48^ We examined inhibition of 1,000 input WT and M184V HIV-RT (10 µL sample) using ISL-TP (**Figure 7B**) and distinguished between WT and resistant samples at 0.5 to 25 µM ISL-TP using post-hoc multiple comparisons test (**Table S7**).

When testing other HIV-RT mutations, we distinguished between 1,000 WT and 1,000 K65R HIV-RT (10 µL sample) using TFV-DP at drug concentrations ≥ 25 µM (**Figure 7C, Table S7**), and between 1,000 WT and 1,000 Y188L HIV-RT (10 µL sample) using the NNRTI DOR at drug concentrations ≥ 0.5 µM (**Figure 7D, Table S7**). K65R and Y188L mutations confer >2 and 50- fold resistance to TFV and DOR, respectively.^49–51^ For testing with ISL-TP and TFV-DP, which are dATP analog drugs, we decreased dATP to 50 µM while keeping remaining dNTPs at 200 µM; since DOR is a non-analog drug (NNRTI), we kept all dNTPs equal at 200 µM. Raw qPCR curves and inhibition curves using Ct values are shown in **Supplementary Figures 10** and **11**, respectively.

### Assay Variation and Cost Analysis

To assess the reproducibility of the PERT assay results, we compared intra- and inter-assay variation. While most experiments were run using 1 cDNA replicate divided into triplicates for qPCR, for analysis of intra-assay variation we instead ran 3 cDNA replicates, each divided into duplicates for qPCR (total N = 6, **Figure 8A**). We ran a standard curve to assess variation across HIV-RT input quantities; one NRT sample was detected (5 NRT undetected and all 3 NTC undetected) while all HIV-RT samples were detected. All coefficient of variation (CV) values were < 3% for samples containing HIV-RT (3.24% CV for NRT samples) (**Table S8**). Raw qPCR curves for the intra-assay experiment can be seen in **Supplementary Figure 12**.

Inter-assay variation was assessed by combining optimized PERT results over > 6 months, two CFX-96 machines, varied sample volumes (1 or 10 µL sample), multiple changes in reagent stocks, and changes in temperature/humidity due to change in season. We compared the Ct values for detecting each input HIV-RT quantity from 10 to 10^6^ (VL testing format, **Figure 8B**) and CVs remained < 5%, including for NRT samples (**Table S8**). While 12% (15/126) of NRT samples were detectable within 45 cycles of qPCR, the average of NRT samples remained well- above that of the lowest HIV-RT input quantity (42.13 for NRT versus 38.14 for 10 HIV-RT) and 0 NTC samples were detected, with off-target detection therefore not posing a concern for false- positive samples. In future validation with clinical samples, we will set a cycle cut-off to distinguish NRT controls from low VL samples and will use probit analysis^52^ to determine the analytical limit of detection. Additionally, since some lots of RNasin may have low levels of RT activity, each order will be evaluated prior to use.^22^ Finally, to assess inter-assay variation for the DRT assay format, we compared the detection of 1,000 input HIV-RT at the 0 drug condition across varied dNTP concentrations: all equal dNTP or with decreased nucleotide that a drug is an analog for (while keeping the remaining dNTPs equal, **Figure 8C**). Again CVs remained ∼3% (**Table S8**) and dNTP concentrations did not have a significant impact on assay variability.

To provide a brief cost analysis of the current iteration of the PERT assay (with reagents purchased for lab use), we estimated the cost of running each cDNA synthesis replicate and qPCR amplification replicate, with the 10 and 20 µL cDNA synthesis reaction formats and 3TC- TP at 0 or 500 µM (**Table S9**). Based on our calculations, running one cDNA synthesis replicate costs $0.72–1.77 (USD) and running one qPCR amplification replicate costs $0.49, excluding consumables such as tubes, Kimwipes™, and PCR plates. Additional requirements within the WHO TPP call for an assay cost of < US $40 (ideally < US $10), which should readily support a variety of combinations of cDNA and qPCR replicates, particularly with larger-scale purchasing.

### Outlook and future work

We anticipate that the need for accessible DRT for the variety of HIV drug classes that are affected by HIV-RT mutations will continue to increase with funding cuts and with the continued rollout of new classes and formulations of ART,^53,54^ including the newly approved once-daily oral regimen of DOR in combination with ISL. As of 2023, an estimated 93% of all PLWH in LMICs were taking ART regimens containing dolutegravir,^55^ an INSTI with a high genetic barrier to resistance. Although the first-line tenofovir, lamivudine, and dolutegravir (TLD) regimen can remain effective despite NRTI-drug resistance mutations,^56^ studies have shown an association between underlying NRTI resistance and eventual acquisition of dolutegravir resistance.^57–59^ In its current iteration, the PERT assay can only detect resistance to RT inhibitors, but could still inform risk for eventual treatment failure with dolutegravir. Thus, more accessible RT inhibitor DRT with PERT may help to increase the longevity of TLD regimens and prevent increased dolutegravir resistance.

In addition, long-acting injectable (LAI) formulations of RT inhibitor drugs are now available for HIV treatment and have been shown to produce high rates of viral suppression, even in populations with prior viremia and/or adherence challenges.^60^ The NNRTI rilpivirine is included in FDA-approved bi-monthly LAI HIV treatments along with the INSTI cabotegravir.^61^ However, LAIs are not recommended for PLWH who have resistance to either drug in the formulation, and consequently, assays like PERT could support the tailored rollout of rilpivirine and other emerging long-acting RT inhibitors in real-world settings.

We acknowledge a limitation of our current work is the absence of a comparison of the viral load and drug resistance measurement of this assay to other gold-standard methodologies with matched or paired specimens. Other than the proof-of-concept testing of HIV-RT activity detection from cell culture supernatant, we used recombinant HIV-RT enzyme for all testing and therefore require validation of VL measurements and DRT from plasma clinical samples.

Additional limitations include the requirement of cold-chain storage, reagent stability, and machinery capable of precise thermocycling (∼ US $12,000-40,000). While the current minimal WHO TPP guidelines support the use of each of these, reduced reliance on external equipment/infrastructure would enable more widespread impact in point of care settings. Our goal in this manuscript was to demonstrate the feasibility in meeting WHO TPP requirements for viral load and drug resistance testing and our ongoing work is focused on sample preparation and assay validation.

## CONCLUSIONS

We optimized a phenotypic enzymatic assay for quantifying HIV viral load and identifying drug resistance based on the cDNA synthesis activity of HIV-RT enzyme in the presence or absence of RT inhibitor drugs followed by qPCR readout. Due to the well-established accuracy of RT-qPCR and the capability of enzymatic assays to detect resistance arising from a wide variety of mutations, this assay has the potential to provide sensitive and actionable information about the efficacy of HIV treatment using readily available instruments in most clinical laboratories.

We optimized various assay parameters, including RT reaction temperature, dNTP concentration, and qPCR primer and probe concentrations to enable detection of low levels of HIV-RT enzyme—as low as 10 molecules (corresponding to a VL of ∼10 RNA copies/mL). We then showed initial proof of extraction-free HIV-RT activity detection using a modified buffer supporting direct lysis of HIV virions and the presence of plasma with minimal inhibition.

Compared to other enzymatic assays for measuring HIV-RT activity, we have the highest sensitivity (detection of the lowest number of HIV-RT molecules and lowest HIV VL) and one of the shortest total assay times (∼2 hours). Furthermore, we showed that this PERT assay could differentiate specifically between WT and resistant HIV-RT in the presence of 3TC-TP, ISL-TP, TFV-DP, and DOR. Next, we showed that PERT could distinguish between WT and mixed populations of WT and M184V HIV-RT, which is more representative of real-world drug resistance. This assay could detect as little as 10% mutant fraction at 1,000 HIV-RT (VL ∼2,500) when comparing Ct values at 0 µM and 500 µM 3TC-TP. Based on the assay’s ability to detect low levels of HIV and resistance, it may be a useful tool for estimating both VL (at 0 µM 3TC-TP) and drug resistance (at 500 µM 3TC-TP), thereby reducing the required number and cost of tests for monitoring HIV treatment. Given the critical and growing need for HIV DRT we believe there is a need for more accessible diagnostic technologies that can provide actionable information in a variety of real-world settings. This work lays the foundation for revisiting PERT as a tool for monitoring the HIV epidemic in local and global health contexts.

## Supporting information

Mims et al PERT Supplementary File

## Author Contributions

**Dorothy Mims:** Conceptualization; Formal Analysis; Investigation; Methodology; Writing – Original Draft Preparation; Writing – Reviewing and Editing; Visualization; Project Administration. **Megan Chang:** Conceptualization; Funding Acquisition; Methodology; Writing – Reviewing and Editing; Visualization. **Ayokunle Olanrewaju:** Conceptualization; Funding Acquisition; Methodology; Project Administration; Resources; Writing – Reviewing and Editing.

## Acknowledgements

The authors are grateful for funding the Washington Entrepreneurial Research Evaluation and Commercialization Hub (WE-REACH), from the Arnold and Mabel Beckman Foundation, and from the University of Washington STD AIDS Training Grant (T32 AI07140). We are also grateful for helpful input and resources from Dr. Qin Wang, Dr. Barry Lutz, Enos Kline, Dr. Geoffrey S. Gottlieb, and Robert A. Smith.

## Data Availability

Raw and analyzed data are available on Zenodo.

