## Supplementary material for "An Optimized Product-Enhanced Reverse Transcriptase Assay for Sensitive and Quantitative Detection of HIV Viral Load and Phenotypic Drug Resistance": Mims et al PERT Supplementary File

**Author Affiliations:**

Linear regression equations and corresponding coefficients of determination (R^2^) and qPCR efficiencies (with 95% confidence intervals) for each condition tested in the optimization and volume scale-up experiments for the PERT assay are shown in Supplementary Table 1. Efficiency of each qPCR reaction was determined as follows:

$Efficiency=100 x \left( -1+{10}^{\left( -1/\mathrm{slope} \right)} \right)$	(II)

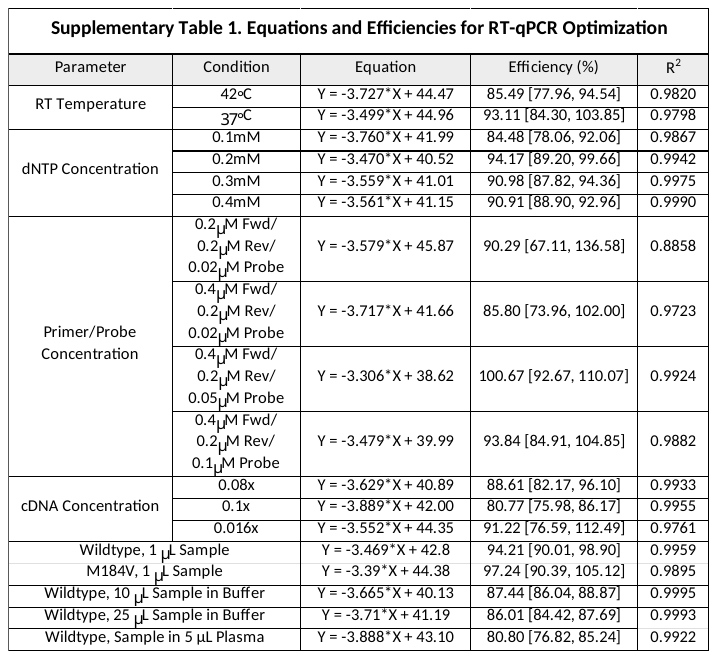

Calculations for determining the ratio of each deoxynucleotide (dNTP) to RNA template for various dNTP concentrations are shown in Supplementary Table 2.

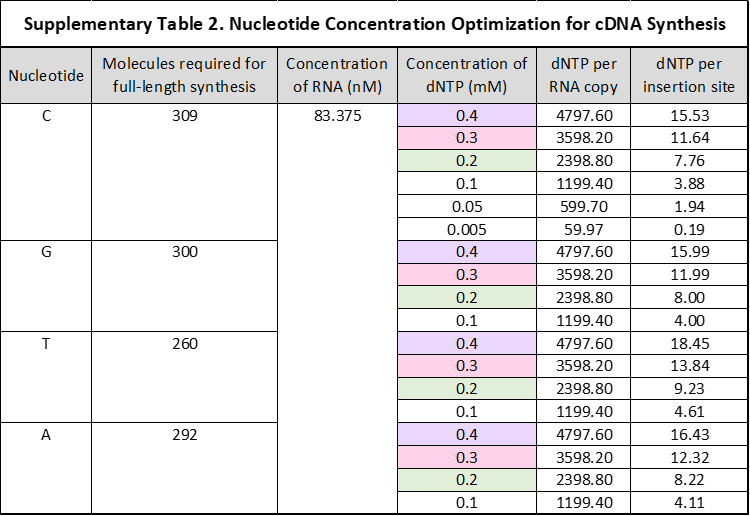

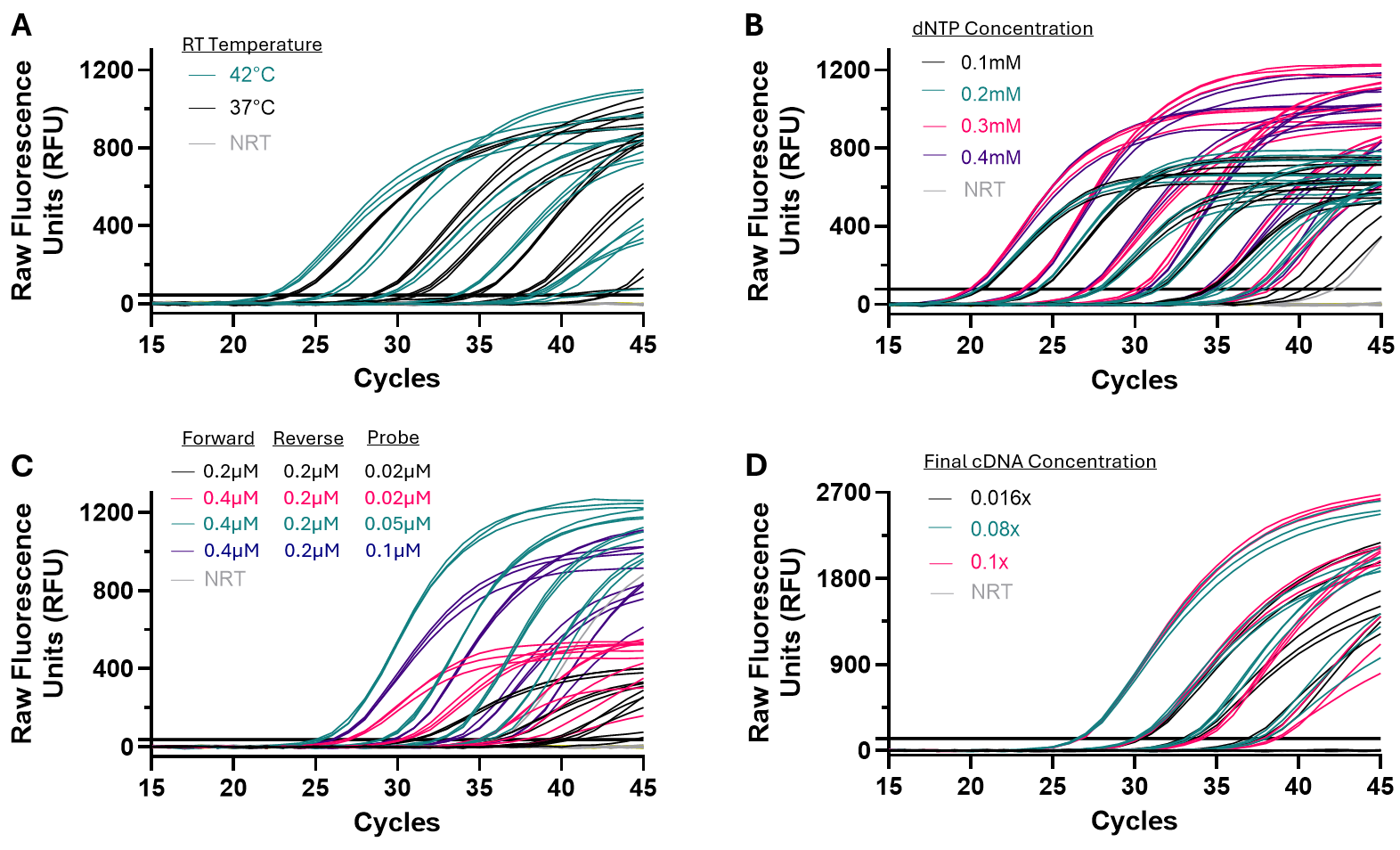

**Supplementary Figure 1. Raw qPCR curves** **for optimization of the PERT assay at each condition: (A)** RT reaction (cDNA synthesis) at 37°C and 42°C, **(B)** varied dNTP (dGTP, dATP, dCTP, and dTTP) concentration in the RT reaction, **(C)** qPCR primer and probe concentration, and **(D)** cDNA concentration in qPCR reactions. Optimal conditions for each experiment are shown in green. Experiments were completed by dividing each completed cDNA sample into three technical qPCR replicates (N=3). NRT controls are also shown, but NTC controls were not detected.

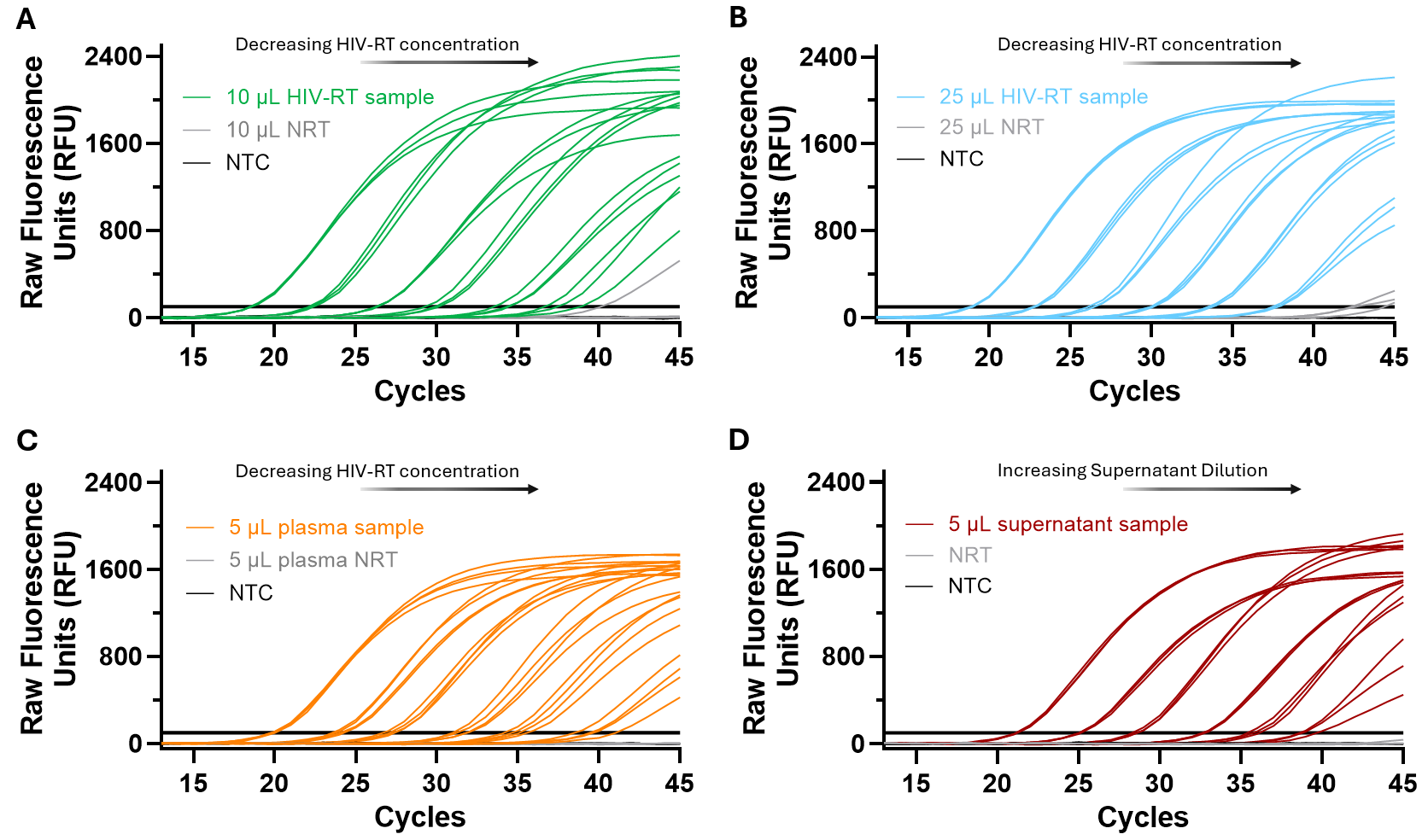

**Supplementary Figure 2. Raw qPCR curves** **for detection of HIV-RT activity using larger sample volumes, plasma, and culture supernatant. (A,B)** Increasing cDNA synthesis reactions to 20 and 35 µL allows scale up of HIV-RT sample volume to 10 and 25 µL, respectively. **(C)** Spiking 1 µL HIV-RT into 5 µL HIV-negative plasma allows amplification despite the presence of inhibitors from plasma samples. **(D)** Using increasing dilutions of HIV-1 cell culture supernatant (diluted in 1X RT Buffer), 5 µL supernatant samples were detected across a range of dilution factors. Experiments for **(A-B)** and **(D)** were completed by dividing each completed cDNA sample into three technical qPCR replicates (N=3), and 2 cDNA replicates of each condition were divided into two qPCR replicates each (N=4) for **(C)**. NRT controls are also shown, but NTC controls were not detected.

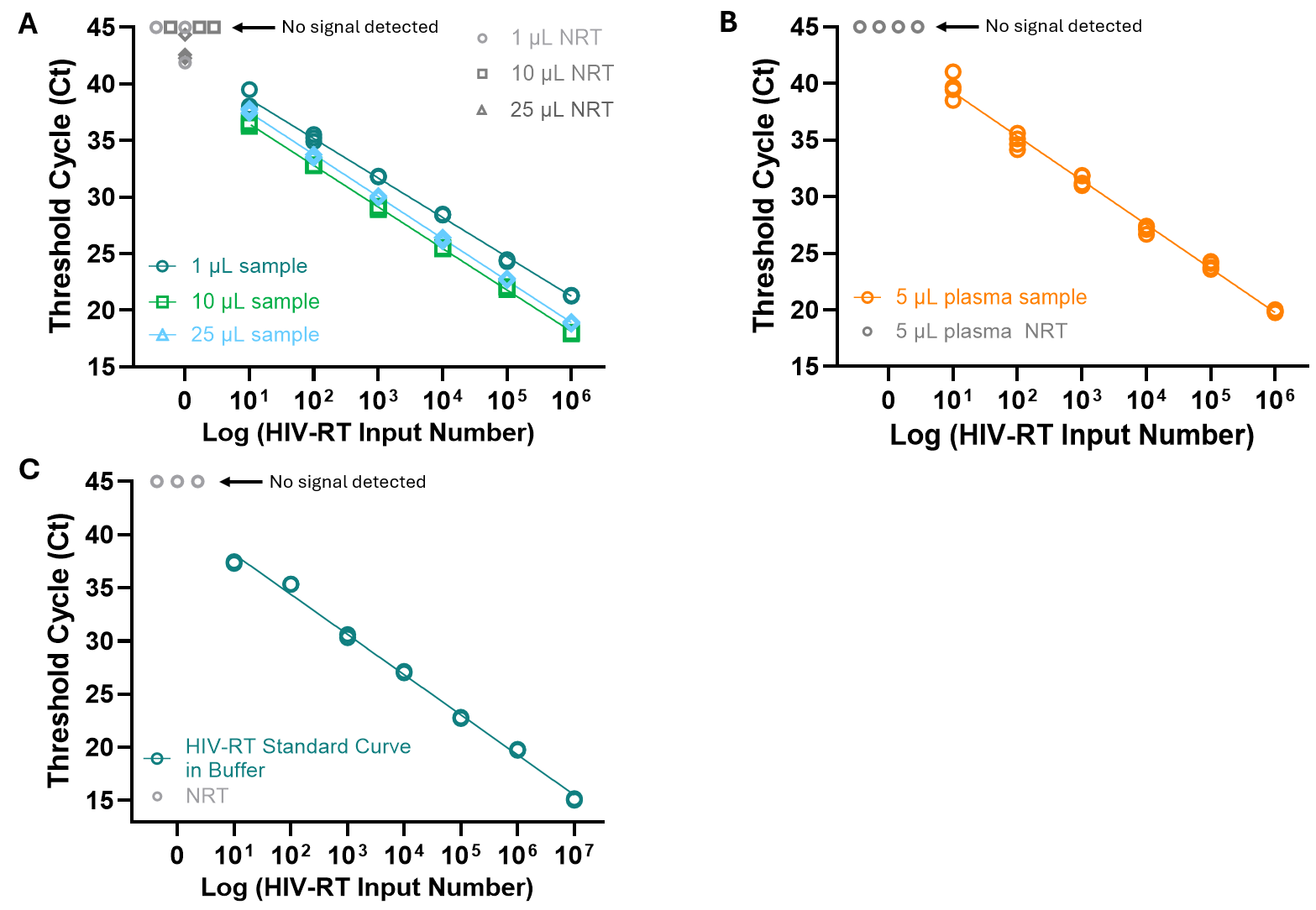

**Supplementary Figure 3. qPCR standard curves** **for detection of HIV-RT activity using larger sample volumes, plasma, and culture supernatant. (A)** Increased cDNA synthesis reactions at 20 and 35 µL with scaled-up HIV-RT sample volume from 1 to 10 and 25 µL, respectively. **(B)** 1 µL HIV-RT spiked into 5 µL HIV-negative plasma allows linear amplification despite the presence of inhibitors from plasma samples, with no detection of NRT plasma samples. **(C)** 10 µL HIV-RT buffer sample standard curve used to determine estimated cell culture supernatant HIV-RT quantity (**Figure 4C**). Experiments for **(A)** and **(C)** were completed by dividing each completed cDNA sample into three technical qPCR replicates (N=3), and 2 cDNA replicates of each condition were divided into two qPCR replicates each (N=4) for **(B)**. NRT controls are also shown, but NTC controls were not detected. Best-fit equations, R^2^ values, and efficiencies for **(A)** and **(B)** can be found in **Table S1**.

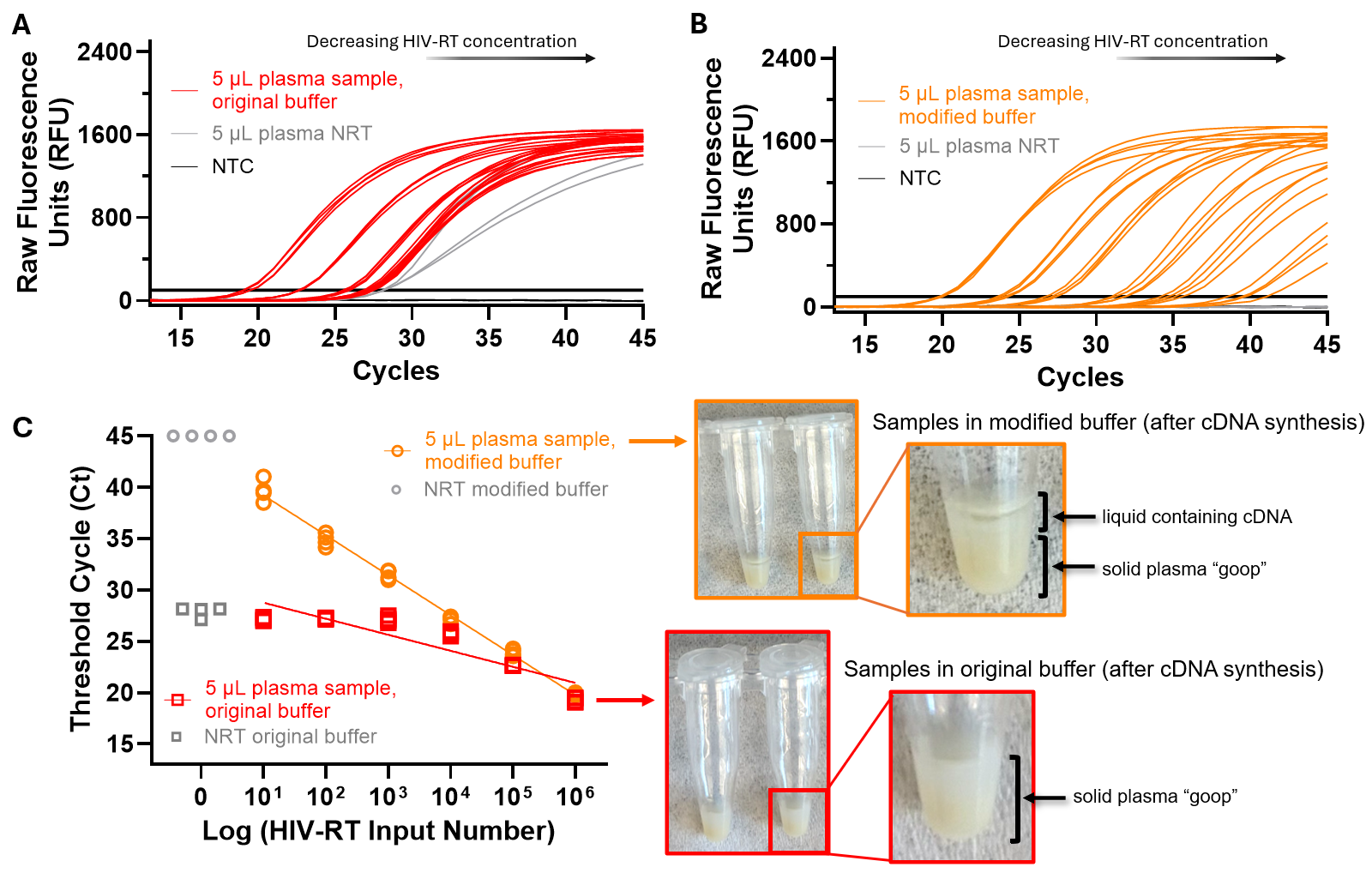

**Supplementary Figure 4. Modified RT Buffer improves the performance of PERT testing using spiked plasma samples. (A,B)** Raw qPCR curves for 1 µL HIV-RT spiked into 5 µL plasma using the original RT buffer (0.16% Triton and 5 mM DTT in 1X Buffer) and modified RT buffer (1.6% Triton and 2.5 mM DTT in 1X Buffer). **(C)** qPCR standard curves for the two buffer formulations, and images showing the separation of liquid from solid plasma “goop” following completed cDNA synthesis. Increased Triton and decreased DTT concentrations allowed greater separation of liquid cDNA from the goop and improved PCR performance. Experiments were completed by dividing each completed cDNA sample replicate into two or three technical qPCR replicates (N=3 and 4 for original and modified buffer). NRT controls are also shown using 1 µL buffer spiked into plasma instead of 1 µL HIV-RT. NTC controls were not detected.

Calculations for determining the viral load (copies HIV RNA/mL) corresponding to each HIV-RT input concentration (HIV-RT molecules and input sample volume) are shown in Supplementary Table 3. We used an estimated 80 HIV-RT/virion for viral load calculations.

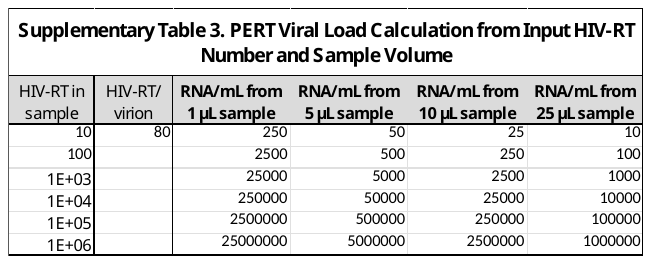

A full comparison of specimen type tested, readout method, sensitivity, and assay run time as well as reference letter (for **Figure 4D**) are shown in Supplementary Table 4.
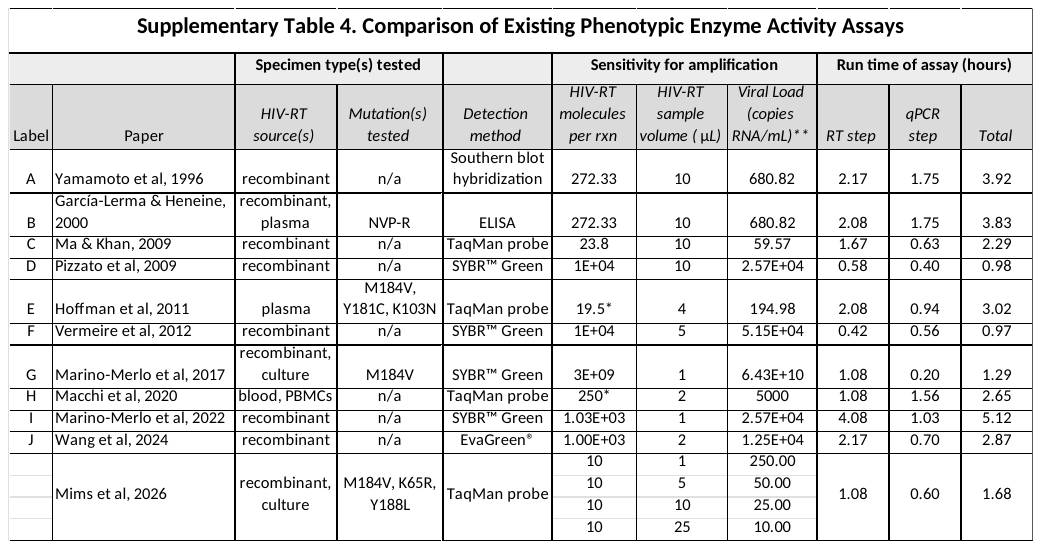

**Sensitivity (molecules HIV-RT per reaction) was calculated from viral load assuming a specific activity of HIV-RT of 5,000U/mg*

***All viral load levels were calculated using the assay’s reported HIV-RT sample volume for cDNA synthesis, assuming 80 molecules HIV-RT/virion and 2 molecules of RNA/virion (except Hoffman et al & Macchi et al, which were reported in terms of viral load).*

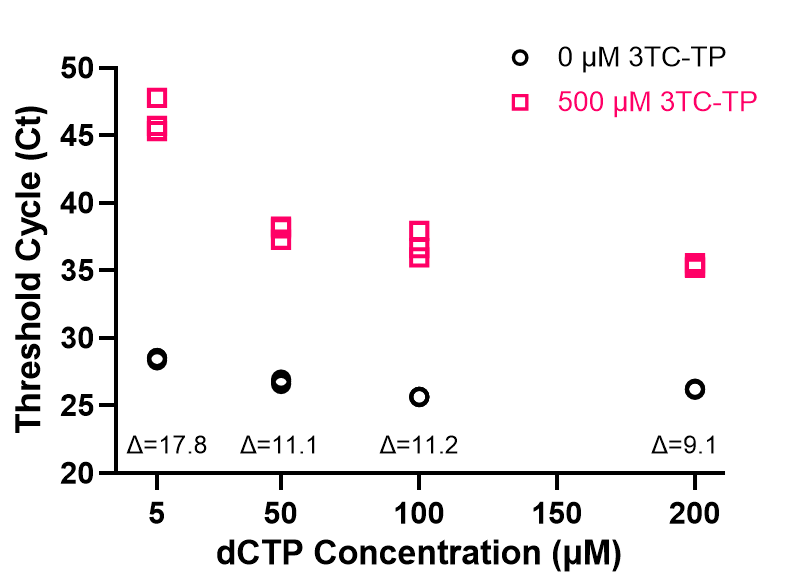

**Supplementary Figure 5. Optimization of nucleotide analog concentration for inhibition of enzyme activity by 3TC-TP.** Ct values for the PERT assay at various dCTP concentrations in the presence of 0 and 500 µM 3TC-TP (10^4^ molecules wildtype HIV-RT). ΔCt values between average drug and no-drug conditions are reported on the graph. Experiments were completed by dividing each completed cDNA sample into triplicates for qPCR (N=3).

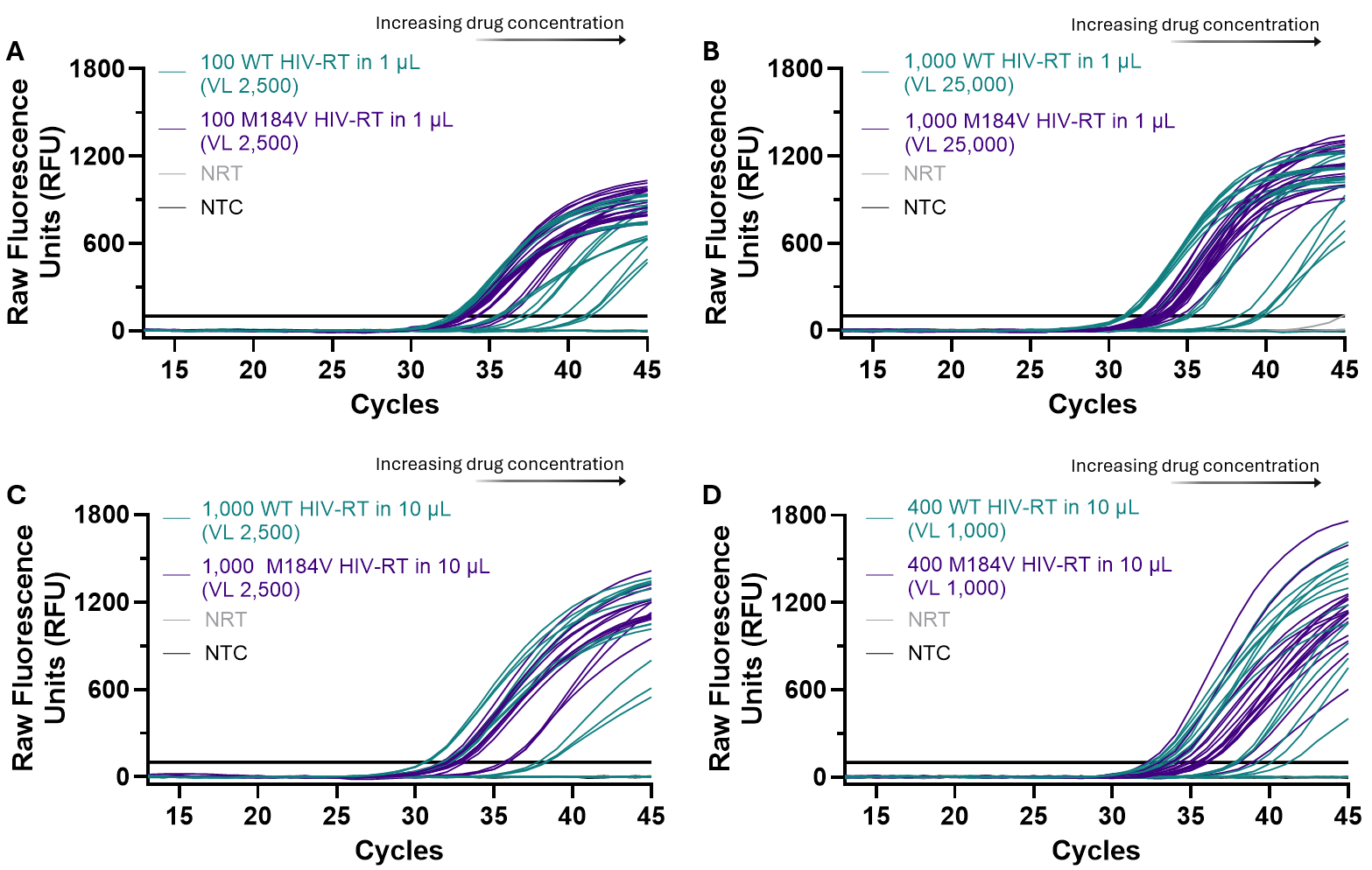

**Supplementary Figure 6. Raw qPCR curves for distinguishing between drug-resistant and drug-susceptible HIV-RT as 3TC-TP concentration increases from 50 or 500 nM to 500 µM. (A)** 100 or **(B)** 1,000 HIV-RT in 1 µL input sample (corresponding to VL 2,500 and 25,000, respectively). **(C)** 1,000 or **(D)** 400 HIV-RT in 10 µL input sample (corresponding to VL 2,500 and 1,000, respectively). Experiments were completed by dividing each completed cDNA sample into three technical qPCR replicates (N=3). NRT and NTC controls are also shown.

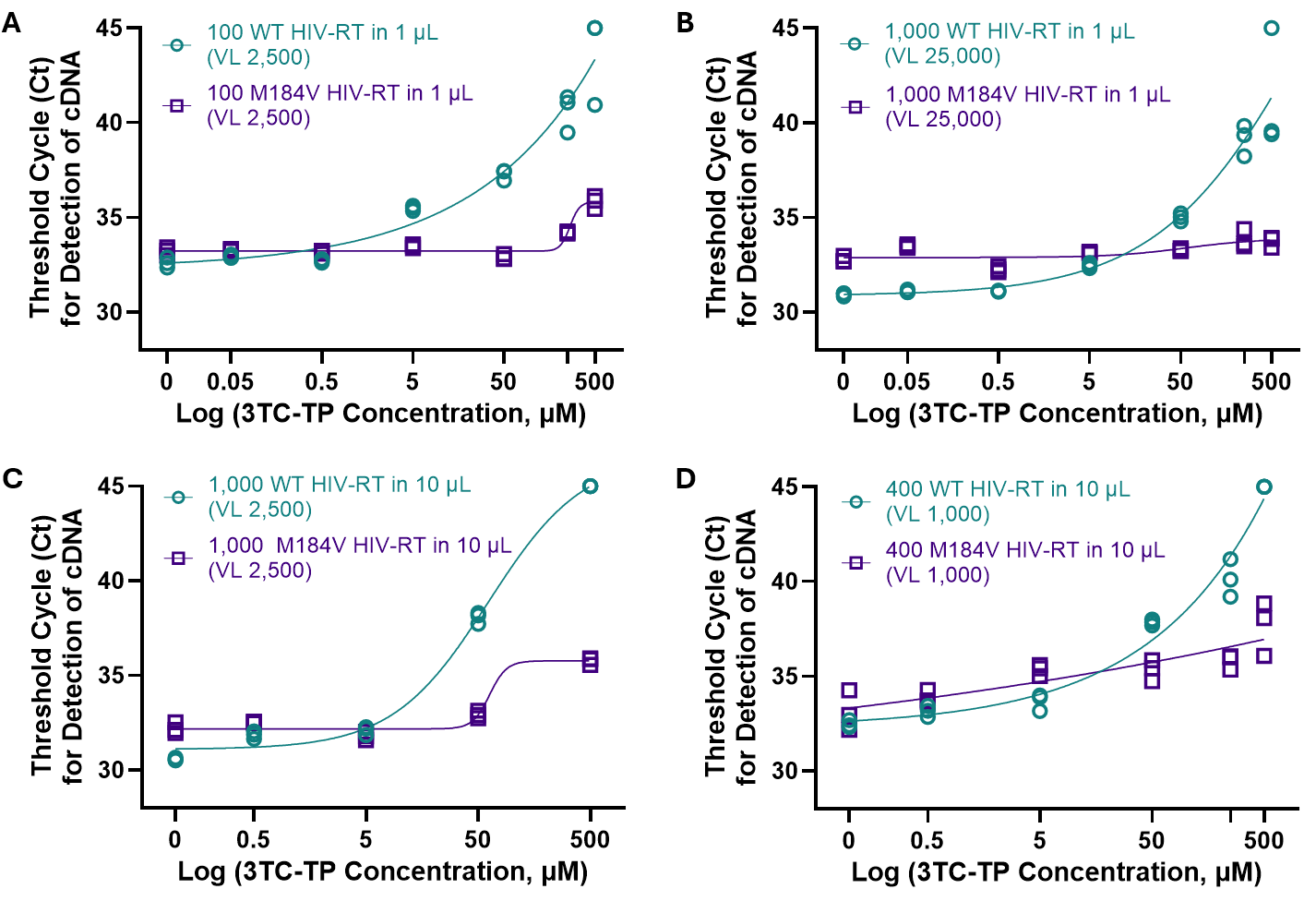

**Supplementary Figure 7.** **Ct values for distinguishing between drug-resistant and drug-susceptible HIV-RT** **as 3TC-TP concentration increases from 50 or 500 nM to 500 µM. (A)** 100 or **(B)** 1,000 HIV-RT in 1 µL input sample (corresponding to VL 2,500 and 25,000, respectively). **(C)** 1,000 or **(D)** 400 HIV-RT in 10 µL input sample (corresponding to VL 2,500 and 1,000, respectively). Experiments were completed by dividing each completed cDNA sample into three technical qPCR replicates (N=3).

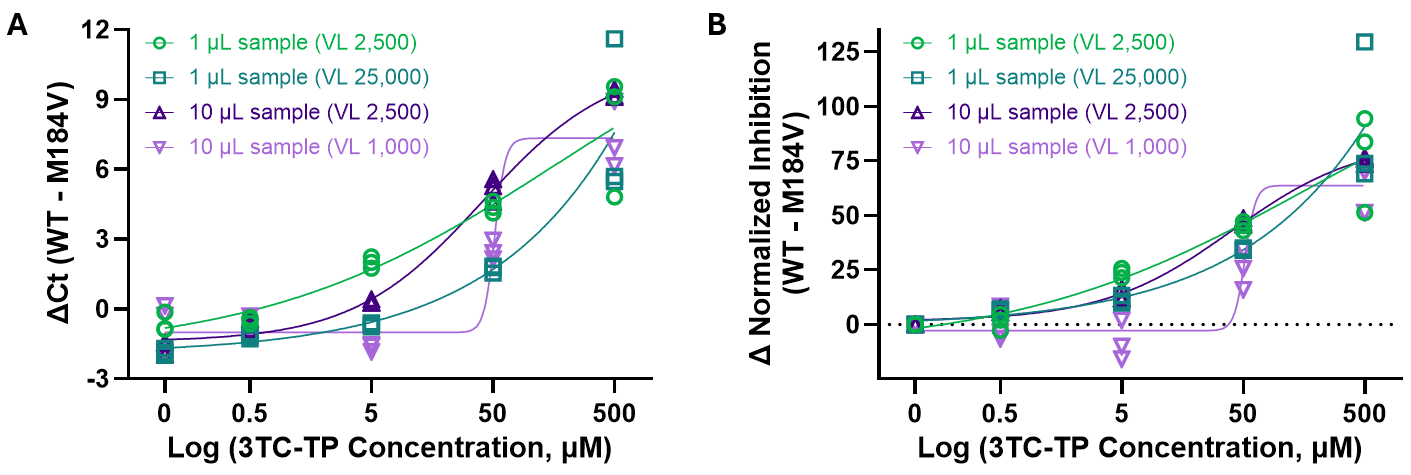

**Supplementary Figure 8. Comparison of ΔCt and normalized inhibition at different input viral loads for 3TC-TP.** For each HIV-RT input concentration, the **(A)** ΔCt and **(B)** difference in normalized inhibition between WT and M184V increases as 3TC-TP concentration increases. Experiments were completed by dividing each completed cDNA sample into three technical qPCR replicates (N=3).

One-way ANOVA followed by multiple comparisons test (Bonferroni) was used to determine statistically significant differentiation between WT and M184V normalized inhibition values at a variety of 3TC-TP and input HIV-RT concentrations (varied total number of HIV-RT molecules and sample volumes). Each inhibition value was normalized to the maximum inhibition condition of WT sample with 500 µM 3TC-TP, with each M184V value compared to the respective WT normalized inhibition value for determination of significance. A full comparison of ΔCt values between drug and no-drug conditions for each WT and M184V sample as well as normalized inhibition values and significance are shown in Supplementary Table 5.

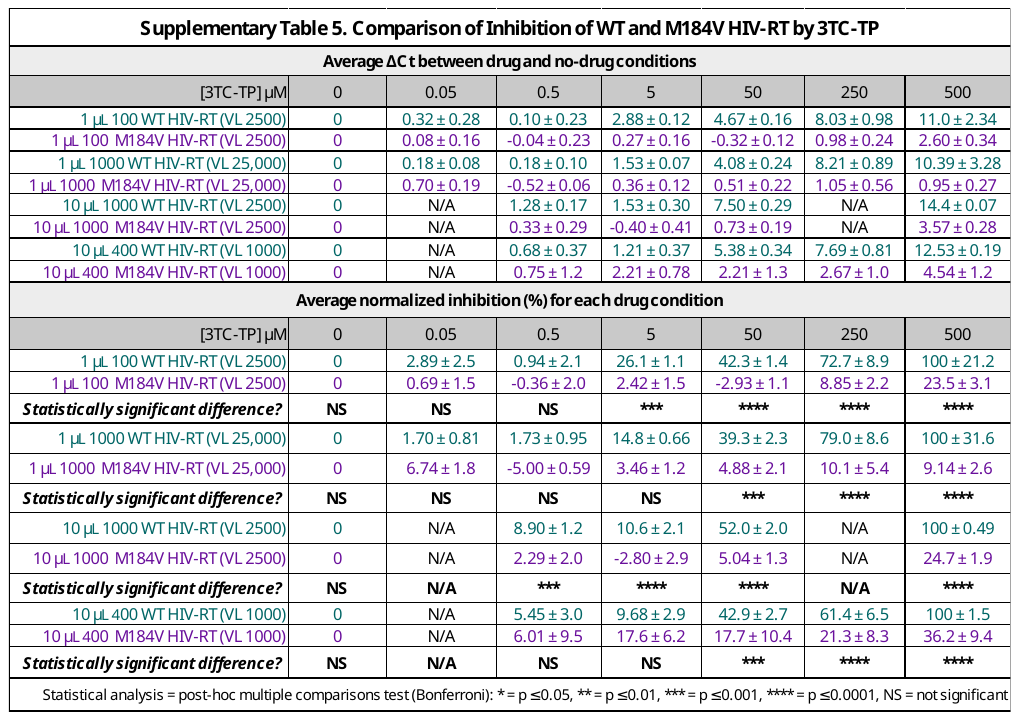

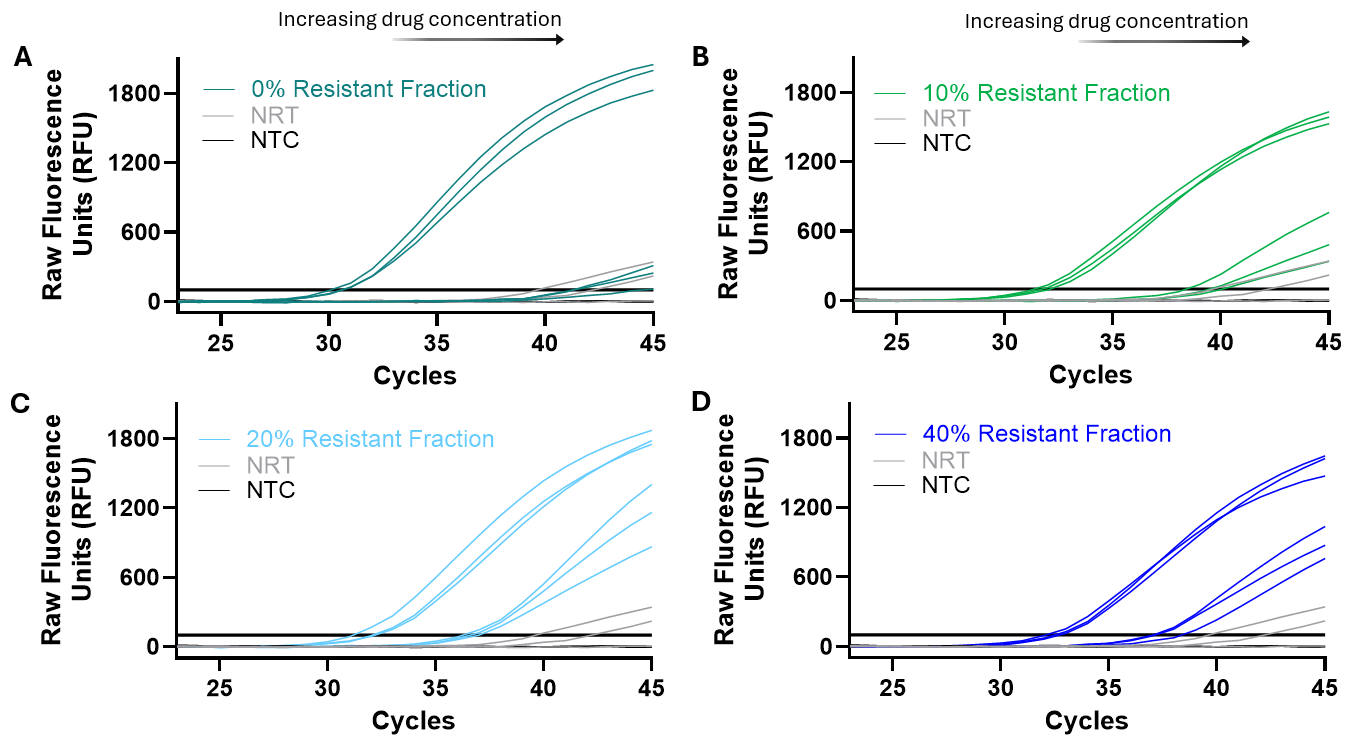

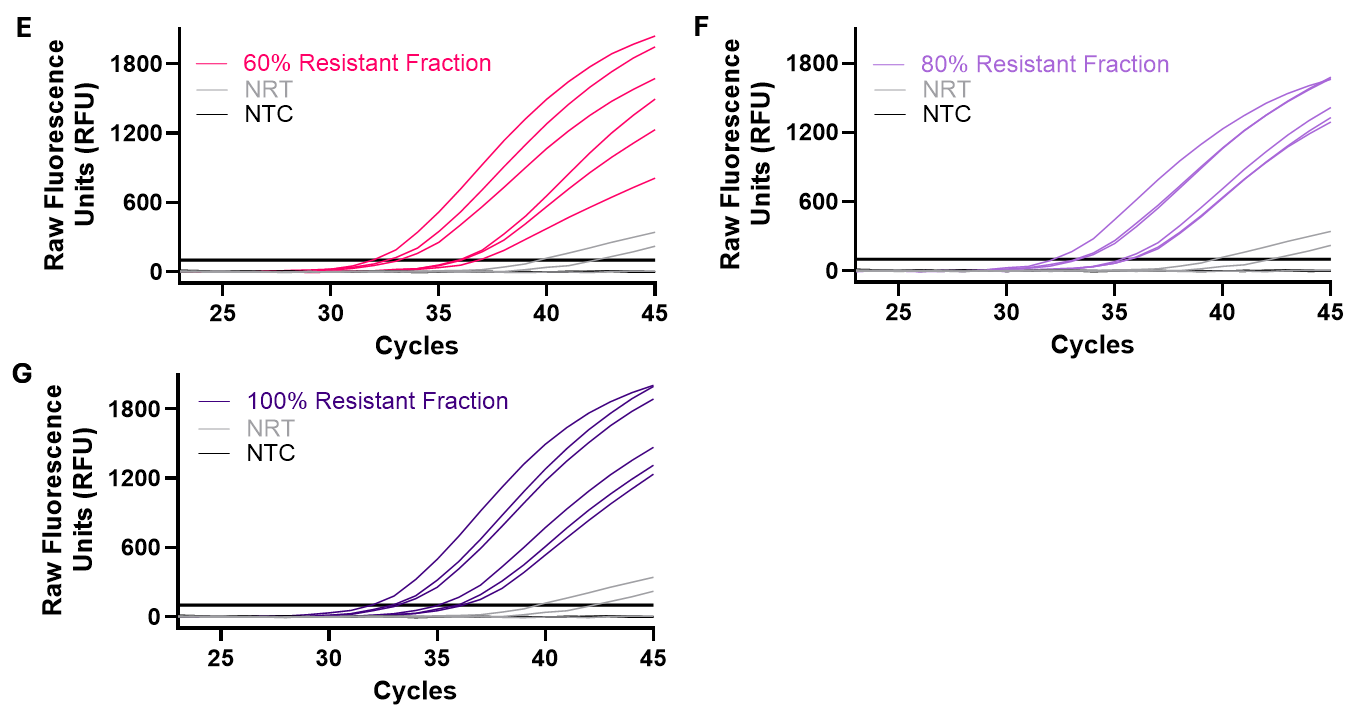

**Supplementary Figure 9. Raw qPCR curves for the PERT assay at 1,000 total HIV-RT molecules with** **(A)** 0%, **(B)** 10%, **(C)** 20%, **(D)** 40%, **(E)** 60%, **(F)** 80%, and **(G)** 100% resistant HIV-RT enzyme fraction at 0 and 500 µM 3TC-TP. NRT and NTC controls are also shown. Experiments were completed by dividing each completed cDNA sample into three technical qPCR replicates (N=3).

One-way ANOVA followed by multiple comparisons test (Dunnett’s) was used to determine statistically significant differentiation of each resistant enzyme mixture from WT (0% resistant) sample. Ct values for each drug condition, ΔCt between drug and no-drug condition, and normalized inhibition values (with statistical analysis) are shown in Supplementary Table 6.

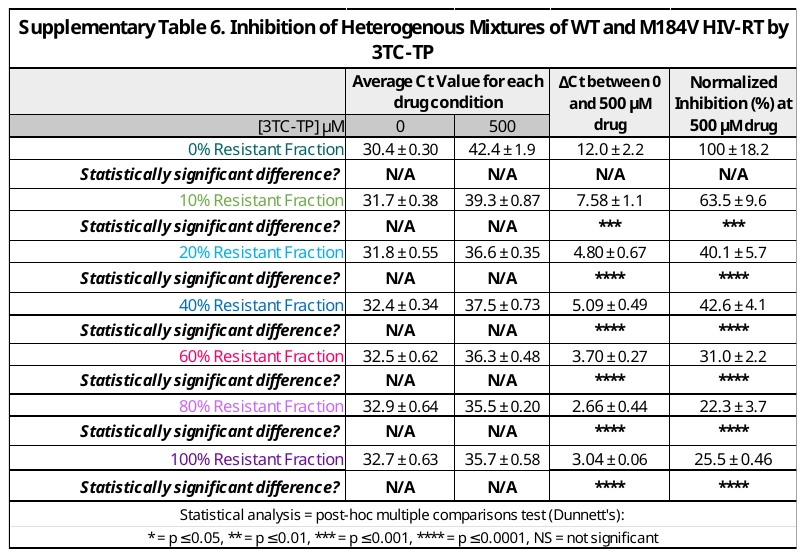

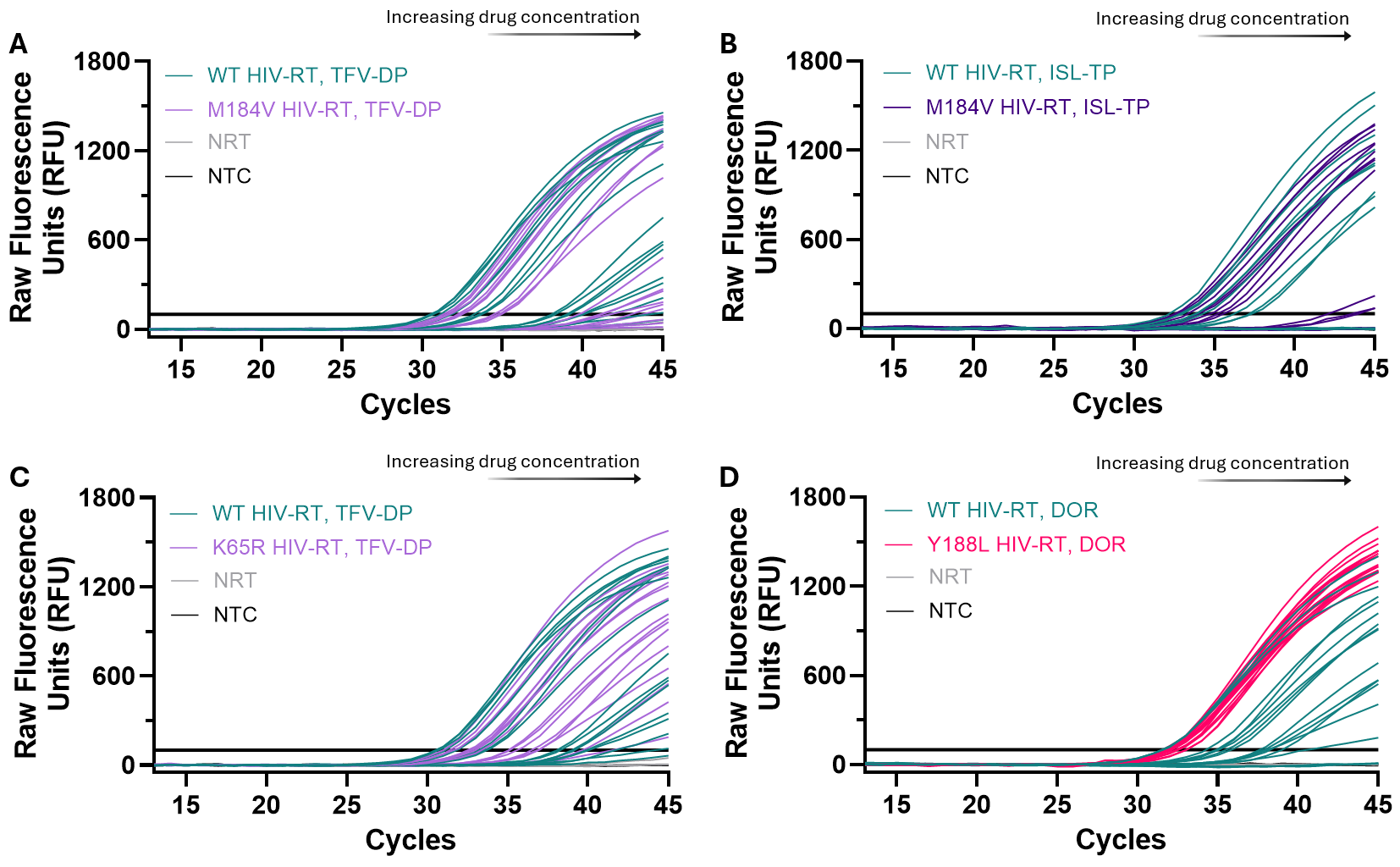

**Supplementary Figure 10. Raw qPCR curves for additional analysis of distinction between drug-resistant and drug-susceptible HIV-RT using (A)** WT and M184V HIV-RT with TFV-DP, **(B)** WT and M184V HIV-RT with ISL-TP, **(C)** WT and K65R HIV-RT with TFV-DP, and **(D)** WT and Y188L HIV-RT with DOR. All experiments used 1,000 WT or mutant HIV-RT in a 10 µL sample volume, and experiments were completed by dividing each completed cDNA sample into three technical qPCR replicates (N=3). NRT and NTC controls are also shown.

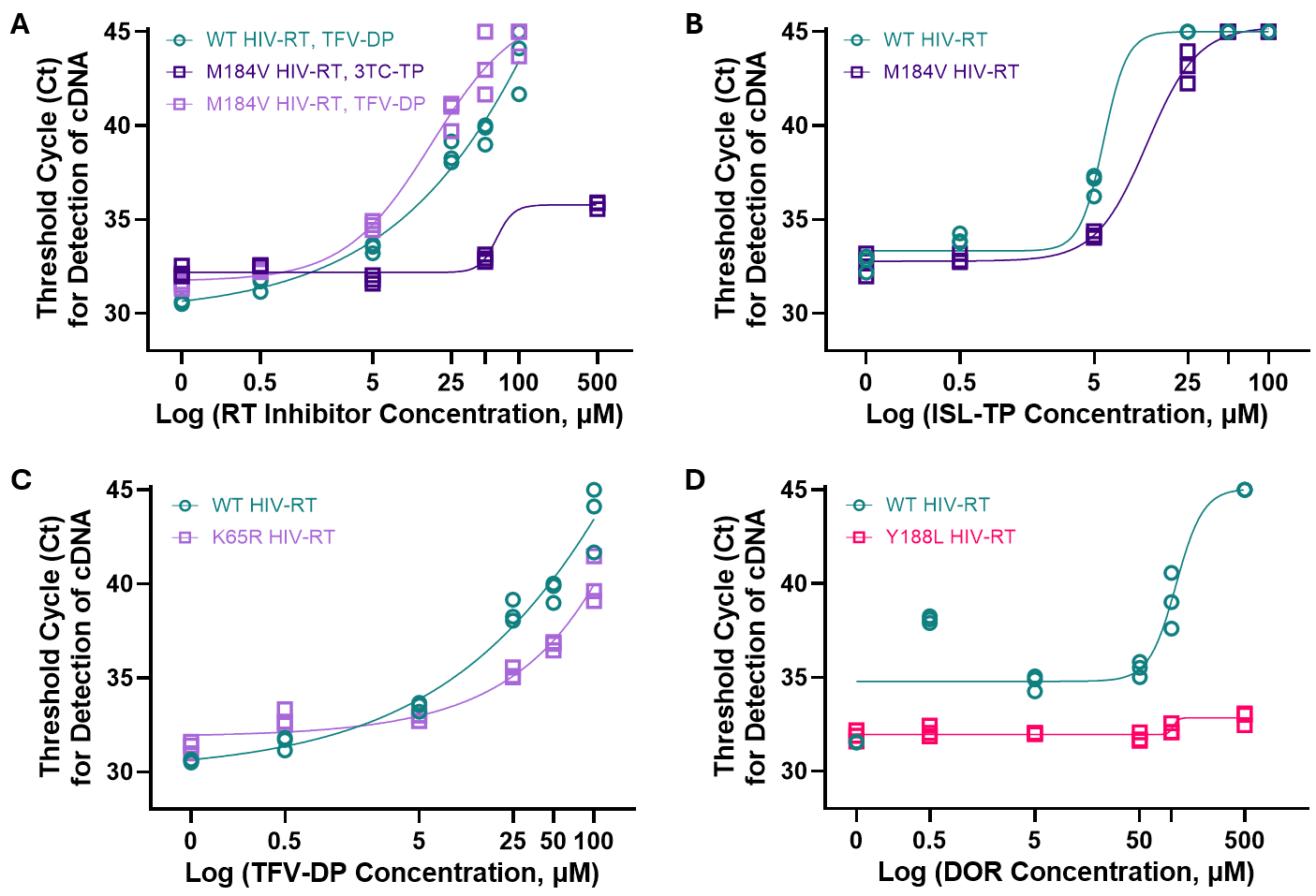

**Supplementary Figure 11. Ct values for additional analysis of distinction between drug-resistant and drug-susceptible HIV-RT using (A)** WT and M184V HIV-RT with TFV-DP and 3TC-TP, **(B)** WT and M184V HIV-RT with ISL-TP, **(C)** WT and K65R HIV-RT with TFV-DP, and **(D)** WT and Y188L HIV-RT with DOR. All experiments used 1,000 WT or mutant HIV-RT in a 10 µL sample volume, and experiments were completed by dividing each completed cDNA sample into three technical qPCR replicates (N=3). NRT and NTC controls are also shown.

One-way ANOVA followed by multiple comparisons test (Bonferroni) was used to determine statistically significant differentiation between WT and mutant HIV-RT normalized inhibition values using a variety of RT inhibitors and concentrations. 1,000 HIV-RT in a 10 µL sample was used for each comparison. Each inhibition value was normalized to the maximum inhibition condition of WT sample with the maximum drug concentration, and each mutant HIV-RT normalized inhibition value compared to the respective WT normalized inhibition value for determination of significance. A full comparison of ΔCt values between drug and no-drug conditions for each WT and mutant sample as well as normalized inhibition values and significance are shown in Supplementary Table 7.

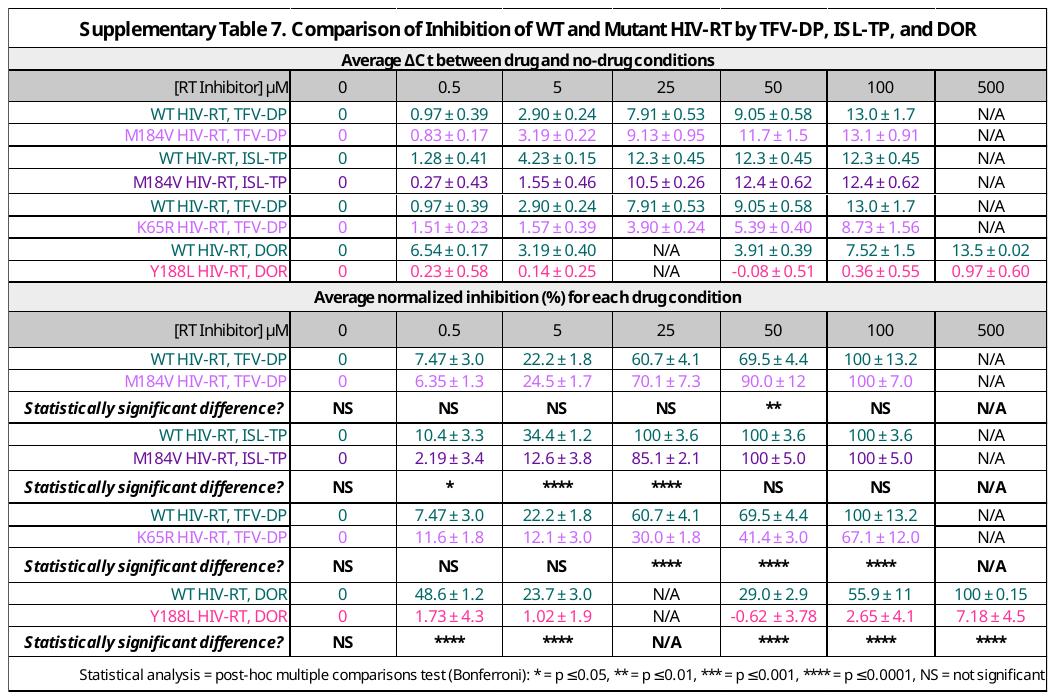

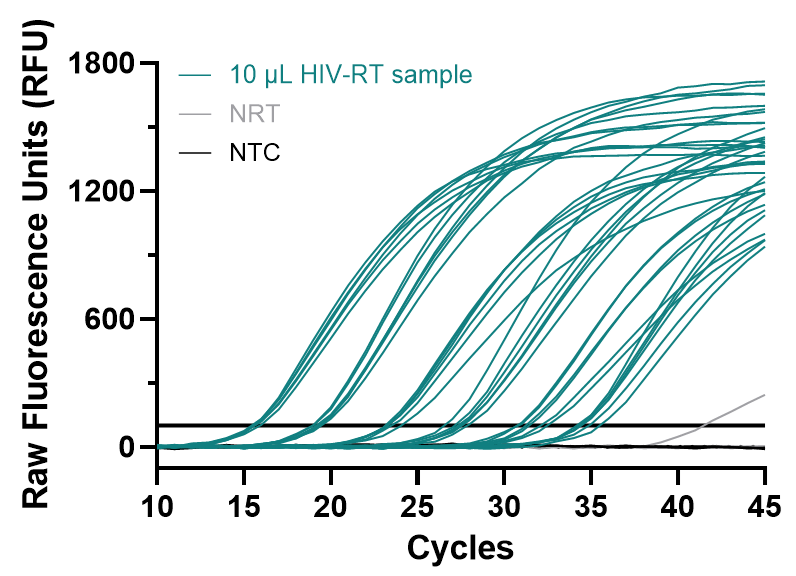

**Supplementary Figure 12. Raw qPCR curves for intra-assay variation experiment.** Three cDNA technical replicates were used for each HIV-RT input concentration (10 µL sample), and completed cDNA samples divided into two qPCR replicates each (total N=6). Three cDNA replicates of NRT and three qPCR replicates of NTC controls were also run and are shown.

A full comparison of intra- and inter-assay variability is shown in Supplementary Table 8. For intra-assay variability, three cDNA replicates of each HIV-RT input concentration were divided into two technical qPCR replicates. For inter-assay variation, we combined assay results over >6 months, two CFX-96 machines, varied sample volumes (1 or 10 µL), and multiple changes in reagent stocks for the viral load measurement and drug resistance testing formats. Coefficients of variation (CVs) are shown for each control and HIV-RT sample.

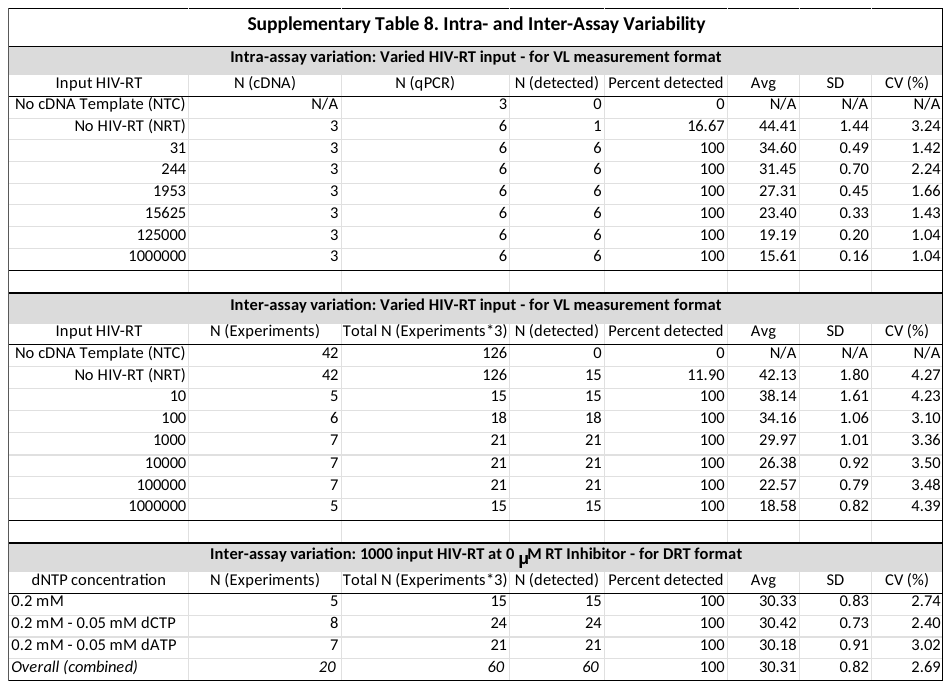

An estimate of the cost to run one cDNA replicate and one qPCR replicate using a 10 or 20 µL cDNA synthesis reaction and 0 or 500 µM 3TC-TP is shown in Supplementary Table 9. This cost estimate excludes consumables used in the assay run.

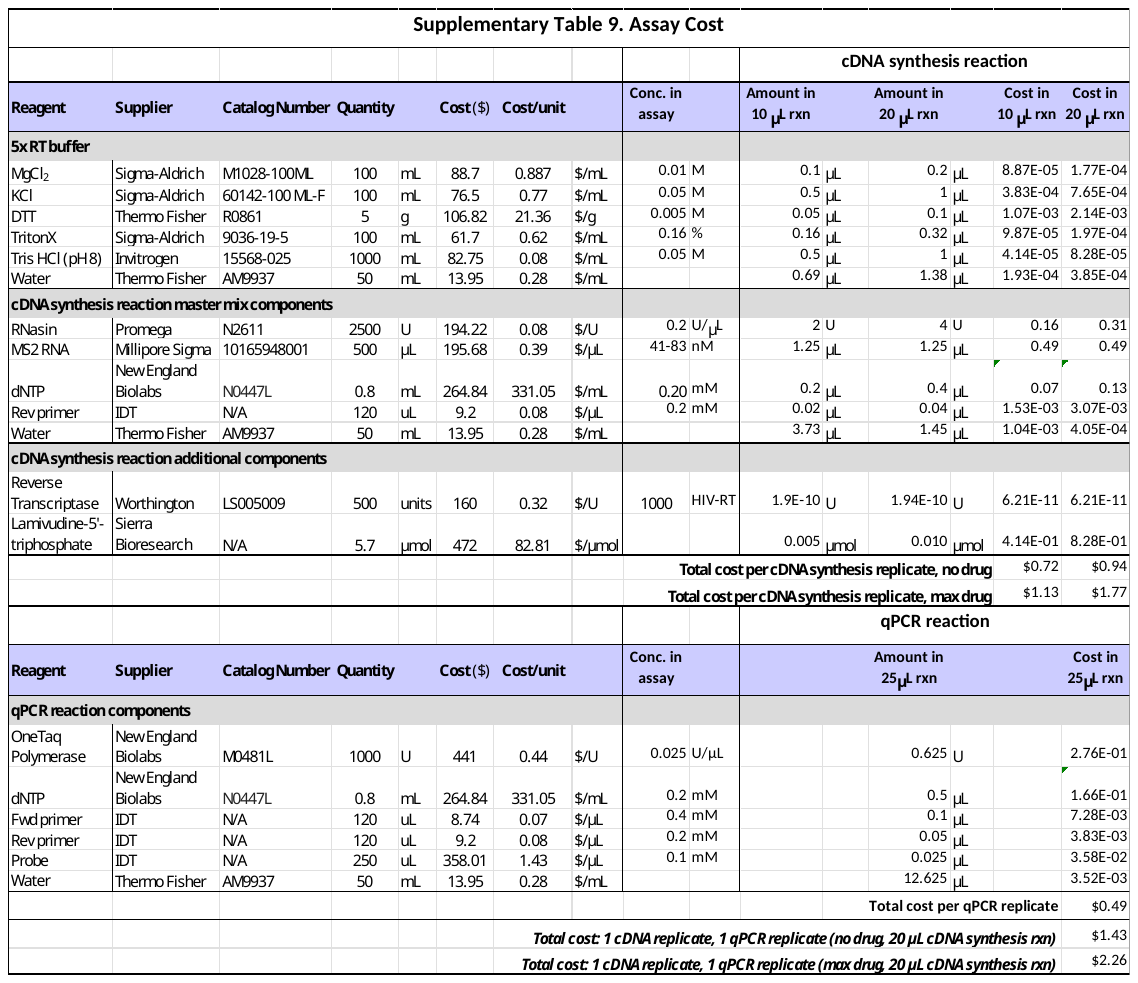

### Full (Detailed) Protocol:

***Step 1. Steps and Reagents for cDNA synthesis (RT) reaction:***

1. Prepare a cDNA Reaction Mix with all components apart from HIV-RT (10µL total reaction volume, with 8-9 µL cDNA Reaction Mix needed per sample). The final concentration of each reagent is listed below.
   - 1x HIV-RT Buffer (from 5x HIV-RT Buffer described above)
   - 0.2 U/µL RNasin® Plus (40U/µL RNasin® Plus Ribonuclease Inhibitor; Promega; Cat. No. N2611)
   - 0.2 mM Deoxynucleotide (dNTP) Solution Mix (10mM dNTP Solution Mix; New England Biolabs; Cat. No. N0447)
   - 0.2 µM Reverse Primer (5’- GCCTTAGCAGTGCCCTGTCT - 3’)
   - 83.375 nM MS2 RNA (667 nM RNA, MS2 from bacteriophage MS2; Millipore Sigma; Cat. No. 10165948001)
   - NF Water (Invitrogen; Cat. No. AM9937) to reach 9 µL volume per sample
2. Pipette 8-9 µL of cDNA Reaction Mix into each PCR tube (clear or white wells).
3. Add HIV-RT (1 µL of sample or 1 µL of 1X RT Buffer for “NRT” control) + 3TC-TP (1 µL per sample).
4. Spin down tubes briefly (<5s) and place on hot plate or in thermocycler.
5. Incubate at 42°C for 60 min + 95°C for 5 min.
6. Dilute resultant cDNA: Add 10 µL nuclease-free water. Vortex tubes or pipette solution a few times to mix, then spin tubes briefly. cDNA is now ready for addition to qPCR reaction mix.

***Step 2. Steps and Reagents for qPCR (amplification and detection) reaction:***

1. (On ice or cold block) prepare a qPCR Reaction Mix with all components apart from cDNA (25 µL total reaction volume; 20 µL qPCR Reaction Mix needed per sample). The final concentration of each reagent is listed below.
   - 0.025 U/µL One*Taq*® Hot Start DNA Polymerase (5,000U/mL; New England Biolabs; Cat. No. M0481)
   - 1x One*Taq* Standard Reaction Buffer (5x One*Taq*® Standard Reaction Buffer Pack; New England Biolabs; Cat. No. B9022 **this buffer comes with M0481**)
   - 0.2 mM Deoxynucleotide (dNTP) Solution Mix (10 mM dNTP Solution Mix; New England Biolabs; Cat. No. N0447)
   - 0.4 µM Forward Primer (5’ - AACATGCTCGAGGGCCTTA - 3’)
   - 0.2 µM Reverse Primer (5’- GCCTTAGCAGTGCCCTGTCT - 3’)
   - 0.05 µM Probe (5’ – FAM – CCCGTGGGATGCTCCTACATGTCA – TAMRA – 3’)
   - NF Water (Invitrogen; Cat. No. AM9937) to reach 25 µL volume per sample
2. Add 20 µL of qPCR Reaction Mix into the bottom of each well of a 96-well PCR plate (Armadillo PCR Plate, 96-well, clear wells; Thermo Fisher Scientific™; Cat. No. AB2396) (or tube if desired).
3. Add 5 µL of cDNA dilution (in three replicates) to the side of each well, ensuring no bubbles are introduced to the reaction. For the “NTC” control, add 5 µL of water instead.
4. Seal plate (Thermo Fisher Scientific™; Cat. No. AB-0558) with even pressure across wells. Ensure tight seal on each well by spraying a Kimwipe™ with 70% ethanol and rubbing plate while applying pressure. Alternatively, use a kitchen scraper tool to evenly scrape and apply pressure over top of entire plate.
5. Briefly (10s) spin down plate and insert in thermocycler.

Start qPCR cycling protocol: 2 min at 95°C followed by 45 cycles of 15s at 95°C + 30s at 56°C.

Discard used qPCR plates and tubes appropriately (Biohazard waste).
